# Functional partnership between carbonic anhydrase and malic enzyme in promoting gluconeogenesis in *Leishmania major*

**DOI:** 10.1101/2020.06.19.161828

**Authors:** Dipon Kumar Mondal, Dhiman Sankar Pal, Mazharul Abbasi, Rupak Datta

## Abstract

*Leishmania* has a remarkable ability to proliferate under widely fluctuating levels of essential nutrients, such as glucose. For this the parasite is heavily dependent on its gluconeogenic machinery. One perplexing aspect of gluconeogenesis in *Leishmania* is the lack of the crucial pyruvate carboxylase (PC) gene. PC-catalyzed conversion of pyruvate to oxaloacetate is a key entry point through which gluconeogenic amino acids are funnelled into this pathway. Absence of PC in *Leishmania* thus raises question about the mechanism of pyruvate entry into the gluconeogenic route. We report here that this task is accomplished in *Leishmania major* through a novel functional partnership between its mitochondrial malic enzyme (LmME) and cytosolic carbonic anhydrase (LmCA1). Using a combination of pharmacological inhibition studies with genetic manipulation, we showed that both these enzymes are necessary in promoting gluconeogenesis and supporting parasite growth under glucose limiting condition. Functional crosstalk between LmME and LmCA1 was evident when it was observed that the growth retardation caused by inhibition of any one of these enzymes could be protected to a significant extent by overexpressing the other enzyme. We also found that while LmCA1 exhibited constitutive expression, LmME protein level was strongly upregulated in low glucose condition. Notably, both LmME and LmCA1 were found to be important for survival of *Leishmania* amastigotes within host macrophages. Taken together, our results indicate that LmCA1 by virtue of its CO_2_ concentrating ability stimulates LmME-catalyzed pyruvate carboxylation, thereby driving gluconeogenesis through pyruvate-malate-oxaloacetate bypass pathway. Additionally, our study establishes LmCA1 and LmME as promising therapeutic targets.

## Introduction

*Leishmania* spp. belongs to the trypanosomatid group of protozoan parasites. They are the causative agents of Leishmaniasis, a poverty-associated neglected tropical disease prevalent in almost 100 countries around the world. With about 12 million affected individuals, an estimated 1 million new cases and 20,000-30,000 deaths every year, leishmaniasis continues to be a global public health problem [1–3]. Depending on the species of *Leishmania* involved, the disease is manifested by a broad range of symptoms that ranges from disfiguring skin lesions to life-threatening infection of the internal organs like liver and spleen [3]. Since *Leishmania* vaccine is still not available, management of the disease solely relies on the limited number of anti-leishmanial drugs. However, wide spread emergence of drug resistant strains, drug-induced toxicity and high cost of treatment highlights the urgency for extensive investigation of unexplored metabolic pathways of the parasite with an eye for novel drug targets [4–7].

One of the fascinating properties of *Leishmania* is its digenetic life cycle, alternating between sand fly vector and mammalian host. During this process, the flagellated promastigote forms of *Leishmania*, that colonizes sand fly midgut, are injected into mammalian host through proboscis. Following this, the parasites are phagocytosed by macrophages, either directly or via apoptotic neutrophils, and are then transformed to non-flagellated amastigotes within the acidic phagolysosomes [8,9]. How *Leishmania* can survive and proliferate in such diverse physiological niche of varying pH and nutrient availability is a fundamental question that has intrigued researchers over the years [10–12]. Metabolic adaptation to fluctuating carbohydrate levels in its surroundings is once such challenging task accomplished by *Leishmania* [11,13].

Hart *et. al.* reported that glucose uptake and utilization in *Leishmania mexicana* promastigotes is several folds higher than in the amastigotes [14]. This is possibly due to the fact that under physiological condition, *Leishmania* promastigotes can easily access glucose from carbohydrate-rich milieu of sand fly midgut [15]. Glucose availability for *Leishmania* amastigotes residing within the phagolysosomal compartment is reported to be much more restricted. Rather, lysosome being the primary site for protein/macromolecular degradation, the environment is rich amino acids and amino sugars [12,16,17]. This change in nutritional environment leads to extensive metabolic reprogramming in the amastigotes that is reflected by significant lowering of glucose transport rate and switch to gluconeogenic mode of energy metabolism whereby intracellular parasites synthesize carbohydrates from non-carbohydrate precursors [11,13]. Although the pathway of gluconeogenesis and its regulation is extensively studied in mammalian system, much less is known about it in lower eukaryotes, particularly in *Leishmania*.

Indispensable role of gluconeogenesis in determining *Leishmania* virulence was first reported by Naderer *et. al.* [18]. By creating a *Leishmania major* null mutant strain of an important gluconeogenic enzyme, fructose-1,6-bisphosphatase (FBP), they showed that the Δ*fbp* mutant amastigotes were unable to grow in cultured macrophage cells or in mice. Interestingly, Δ*fbp L. major* promastigotes, which grew normally in glucose rich medium, were not able to grow at all in glucose depleted medium. The wild type promastigotes could, however, grow in absence of glucose, albeit slowly. Thus, it was evident that gluconeogenesis is also functional in *Leishmania* promastigotes and they may utilize this machinery during occasional period of glucose starvation [18]. Such situation may arise in the sand fly midgut in between two sugar rich meals [15]. Following this initial discovery, glycerol kinase (GK), phosphoenolpyruvate carboxykinase (PEPCK) and pyruvate phosphate dikinase (PPDK) were identified as key players of gluconeogenesis in *L. Mexicana* [19]. Among these enzymes, PEPCK was shown to be upregulated in response to glucose starvation in *Leishmania donovani*, suggesting that the parasite can sense glucose level in its surroundings and accordingly modulate its gluconeogenic activity [20].

Despite these progresses, there are several gaps in our understanding regarding the precise mechanism of gluconeogenesis in *Leishmania*. The mode of pyruvate entry into the gluconeogenic cycle is one such grey area. During prolonged period of glucose starvation in higher eukaryotes, gluconeogenic amino acids, such as alanine, cysteine, glycine, serine, threonine, are first catabolised to pyruvate in the cytosol. Pyruvate is then transported to the mitochondrial matrix with the help of a mitochondrial pyruvate carrier, following which it is converted to oxaloacetate by the enzyme pyruvate carboxylase (PC) [21,22]. This bicarbonate (HCO_3_ ^−^)-requiring carboxylation reaction is a critical step through which the central metabolite pyruvate is channelled into the gluconeogenic pathway [23]. Source of this HCO_3_^−^required for pyruvate carboxylation remained elusive for years. Dodgson *et. al.* provided an important clue by demonstrating that treatment of hepatocytes with carbonic anhydrase (CA) inhibitor, ethoxzolamide, resulted in inhibition of pyruvate carboxylation as well as glucose synthesis in a dose dependent manner [24]. These findings provided the first hint that the mitochondrial carbonic anhydrase V (CAV), by virtue of its CO_2_ hydration activity, might be supplying the crucial HCO_3_^−^ substrate for the PC-catalyzed reaction [24,25]. Unambiguous evidence supporting the functional involvement of mitochondrial CA in gluconeogenesis was provided by Shah *et. al.* by detailed characterization of the CAVA and CAVB knockout mice [26]. Despite these mechanistic insights from the mammalian system (Fig. 1A), how pyruvate carboxylation happens in *Leishmania* is still poorly understood.

**Fig. 1.**
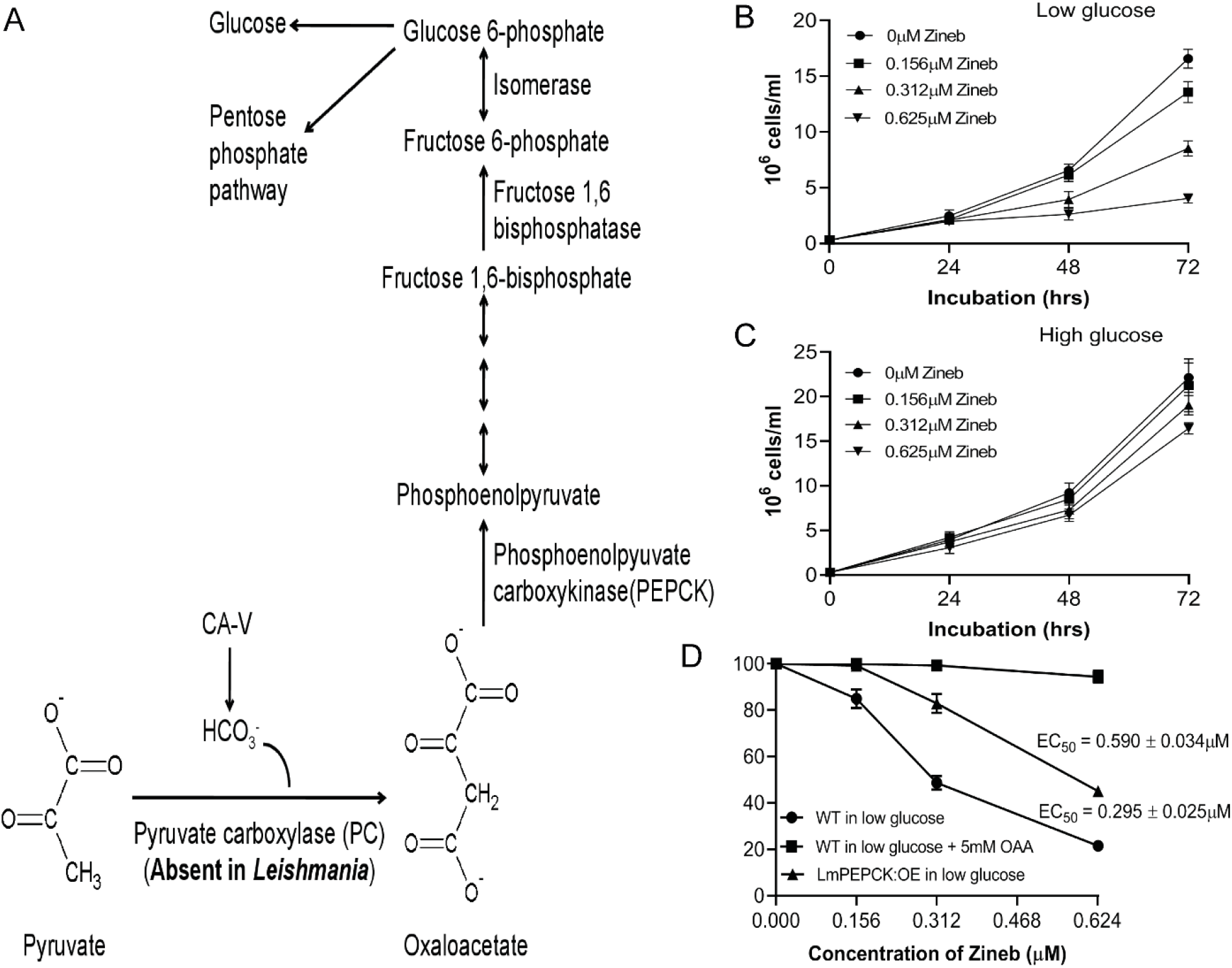
Role of CA in gluconeogenesis in mammalian cells and in *Leishmania*. (A) A schematic representation of the gluconeogenic pathway in mammalian cells showing PC-catalyzed carboxylation of pyruvate to oxaloacetate is dependent upon the HCO_3_^−^ produced by the mitochondrial CAV enzyme. Oxaloacetate is then converted to phosphoenolpyruvate by PEPCK and thereafter gluconeogenesis proceeds through several intermediate steps. It is noteworthy that PC is absent in *Leishmania*. (B, C) Wild type *L. major* promastigotes were grown in low (0.6 mM) or high (6.2 mM) glucose medium (as indicated in the figures) in absence (0μM; circle) or presence of 0.156μM (square), 0.312 μM (triangle) or 0.625 μM (inverted triangle) zineb (CA inhibitor), and growth of the cells was measured by haemocytometer-based cell counting every 24 hrs until 72 hrs of growth. Error bars represent mean ± SD of values from 3 independent experiments. (D) Wild type (WT; circle) or LmPEPCK-overexpressing (LmPEPCK:OE; triangle) promastigotes were grown in low (0.6 mM) glucose medium in absence or presence of indicated concentrations of zineb, and growth of the cells was measured by cell counting after 72 hrs. Zineb-treated WT cells were also grown in presence of 5 mM exogenous oxaloacetate (OAA; square) in low glucose medium. For each experimental condition, cell number in untreated (0 μM zineb) samples was considered as 100%. The EC_50_ values (in μM) of zineb for WT or LmPEPCK:OE *L. major* strain grown in low glucose medium are given in the index. Error bars represent mean ± SD of values from 3 independent experiments.

We recently identified two CAs in *L. major* (LmCA1 and LmCA2) and reported their combined role in maintaining cytosolic pH homeostasis in the parasite [27,28]. Whether they play any other physiological role in *Leishmania*, especially in facilitating the process of pyruvate carboxylation during gluconeogenesis has not been explored so far. In this report, we provide strong evidence to demonstrate that the cytosolic CA isoforms in *L. major* (LmCA1) is functionally involved in promoting gluconeogenesis and in supporting parasite growth under glucose limiting condition. However, exact role of LmCA1 in gluconeogenesis was difficult to explain because of the absence of a bona fide PC gene is in the genome of all *Leishmania* species [29,30]. Also, PC activity could not be detected in *L. mexicana* promastigotes as well as in amastigotes [31]. This led us to the question, does LmCA1 facilitates pyruvate carboxylation via an alternate mechanism? Apart from PC, pyruvate carboxylating activity of malic enzyme (ME), in catalyzing conversion of pyruvate to malate, has been previously reported in few cases in mammalian cells as well as in *Arabidopsis thaliana* [32–36]. Furthermore, presence of a functionally active ME from *L. major* (henceforth referred to as LmME) has recently been identified. Although the kinetic parameters of this enzyme have been determined, its role in *Leishmania* physiology is yet to be deciphered [37]. We were thus provoked to hypothesize that LmCA1 might be functionally cooperating with LmME in facilitating pyruvate carboxylation and in driving gluconeogenesis through pyruvate - malate - oxaloacetate bypass pathway. We tested this hypothesis by a combination of pharmacological inhibition and genetic overexpression studies and showed for the first time that this enzyme indeed has a pyruvate carboxylating activity and plays an important role in gluconeogenesis in cooperation with LmCA1. We further demonstrated that LmME is localized in the mitochondria and its expression is upregulated under glucose limiting condition. Finally, by performing macrophage infection experiments it was proven that both LmME as well as LmCA1 are required for intracellular survival of *Leishmania*. Collectively, these results resolved an important paradox with respect to pyruvate carboxylation in *Leishmania* thus helped in better understanding of its gluconeogenic pathway.

## Results

### Treatment with CA inhibitor caused *L. major* growth inhibition under glucose-limiting condition due to reduced gluconeogenesis and ATP production

Prior studies have implicated role of mammalian CAV, localized in mitochondria, in synthesizing glucose from pyruvate. It was suggested that CAV promotes gluconeogenesis by facilitating the bicarbonate-dependent carboxylation reaction catalyzed by pyruvate carboxylase (Fig. 1A) [24–26]. Although genome of all *Leishmania* species lack evidence for the presence of a bona fide pyruvate carboxylase gene, we were curious to check if one of the LmCAs may still participate in the gluconeogenic process [28,29]. For this, we grew *L. major* promastigotes in absence or presence of increasing concentrations of zineb, which was recently identified as a potent inhibitor of CA activity in *L. major* [27]. Interestingly, zineb caused a dose-depended inhibition of parasite growth when they were cultured in low glucose medium (0.6mM glucose) in presence of several gluconeogenic amino acids. But this treatment had a minimal effect when 5.6mM exogenous glucose was added to the growth medium (Fig. 1B, C). These results suggested that CA activity might be crucial for *Leishmania* to synthesize glucose from non-carbohydrate precursors. Involvement of LmCA in gluconeogenesis was further supported by the observation that zineb-mediated inhibition of parasite growth in glucose-limiting condition could be completely prevented by supplementing the growth media with 5mM oxaloacetate, an intermediate of the gluconeogenesis pathway (Fig. 1A, D). To validate this result, we engineered a *L. major* strain overexpressing PEPCK (LmPEPCK:OE), a key upstream enzyme of the gluconeogenic pathway (Fig. S1). Interestingly, LmPEPCK:OE strain was much less susceptible to zineb-mediated growth inhibition (EC_50_ = 0.590μM) as compared to the wild type (EC_50_ = 0.295μM), indicating that overexpression of this gluconeogenic enzyme can mitigate the adverse effect of inhibition of LmCA activity. Finally, we measured glucose and ATP levels in *L. major* cells in absence or presence of zineb (Table 1). It was observed that zineb treatment caused ~50% reduction in the glucose and ATP levels in *L. major* cells growing under glucose-limiting condition (0.6mM glucose). Supplementation of the growth media with exogenous glucose or oxaloacetate provided complete protection against zineb-mediated depletion of glucose and ATP levels thereby providing an unambiguous evidence for the involvement of LmCA in gluconeogenesis in *Leishmania* cells.

**Table 1.** Intracellular glucose and ATP levels of untreated or zineb-treated wild type *L. major* grown in low glucose medium in absence or presence of oxaloacetate or glucose.

| Strains/Treatment<sup>a</sup> | Total intracellular glucose (nmol/10<sup>8</sup>cells)<sup>b</sup> | Total intracellular ATP (nmol/10<sup>8</sup>cells)<sup>c</sup> |
| --- | --- | --- |
| Untreated WT | 11.71 ± 0.62 | 56.67 ± 5.20 |
| WT + 0.625 µM Zineb | 5.96 ± 0.35** | 27.92 ± 2.59** |
| WT + 0.625 µM Zineb + 5 mM OAA | 11.31 ± 0.59 | 56.25 ± 1.52 |
| WT + 0.625 µM Zineb + 5.6 mM Glu | 11.05 ± 0.46 | 55.37 ± 2.25 |

<sup>a</sup>Wild type (WT) *L. major* promastigotes were grown in absence or presence of 0.625 µM zineb in low glucose (0.6 mM) medium for 72 h. Zineb-treated cells were either unsupplemented or supplemented with 5 mM oxaloacetate (OAA) or 5.6 mM glucose (Glu) during the 72 h duration.
<sup>b</sup>Total internal glucose concentration (in nmol) was experimentally determined from 2.5 x 10<sup>8</sup> *L. major* cells grown in low glucose medium.
<sup>c</sup>Total internal ATP concentration (in nmol) was experimentally determined from 8 x 10<sup>6</sup> *L. major* cells grown in low glucose medium.
± indicates SD of values from triplicate experiments.
\*indicates significant difference (\*\*p<0.01) with respect to untreated wild type strain.

### LmCA1 but not LmCA2 is involved in gluconeogenesis in *L. major*

We recently reported that *L. major* expresses a cytosolic and a plasma membrane bound CA and named them as LmCA1 and LmCA2, respectively [28]. So, the obvious question was which of these two LmCAs participate in the gluconeogenesis process? To address this, we utilized the LmCA1^+/−^ and LmCA2^+/−^ heterozygous strains and the corresponding genetic complementation strain of *L. major* that were earlier generated and validated by us [28]. It was observed that in glucose-limiting condition, the LmCA1^+/−^ heterozygous strain grew sluggishly as compared to the wild type strain and exhibited 36% reduction in the total number of cells after 72 hours. The growth rate of the corresponding complementation strain (LmCA1^+/−^:CM) was near-normal thereby confirming that single allele disruption of LmCA1 is indeed responsible for the observed growth defect of the mutant strain in glucose-limiting condition (Fig. 2A). However, the LmCA1^+/−^ strain did not exhibit any growth defect when exogenous glucose was added in the media (Fig. 2B). Exogenous addition of oxaloacetate also could completely restore the growth defect of the LmCA1^+/−^ strain in glucose-limiting condition implying that LmCA1 participates in the gluconeogenesis process (Fig. 2C). It is worth noting that the growth rate of the LmCA2^+/−^strain was similar to that of its wild type counterpart in glucose-limiting as well as glucose rich conditions (Fig. 2A, B).To further test the contribution of LmCA1 and LmCA2 in *Leishmania* cell growth under glucose-limiting condition, we generated two *L. major* strains overexpressing either LmCA1 or LmCA2 (LmCA1:OE, LmCA2:OE) (Fig. S1). The LmCA1:OE, LmCA2:OE and the wild type *L. major* cells were grown in glucose-limiting condition and their susceptibilities to the CA inhibitor zineb were compared. Interestingly, as compared to the wild type *L. major*, cells overexpressing LmCA1 was significantly less susceptible to zineb-mediated growth inhibition (EC_50_ values were0.295μM and 0.597μM for wild type and LmCA1:OE, respectively). LmCA2 overexpressing strain on the other hand were quite similar to its wild type counterpart in terms of zineb susceptibility (Fig. 2D). Collectively, these data suggest that LmCA1, but not LmCA2, plays an important role in sustaining parasite growth in low sugar environment. Apart from sluggish growth phenotype, the LmCA1^+/−^ strain, in glucose-limiting condition, also exhibited crippled morphology with significant reduction in cell length. This phenotype was reversed to a significant extent in the LmCA1^+/−^: CM complementation strain (Fig. 2E, F). Our morphological data suggest that single allele disruption of LmCA1 makes the parasite susceptible metabolic stress in low-glucose environment. Metabolic stress in the LmCA1^+/−^ strain under glucose-deprived condition was even more evident when we found that its glucose and ATP contents was significantly less compared to that in wild type *L. major* (more than 40% drop for both glucose and ATP). As expected, glucose and ATP levels in the LmCA1^+/−^: CM complementation strain was almost same as that in the wild type cells thus confirming that LmCA1 is indeed involved in glucose and ATP synthesis in *L. major*. The LmCA1^+/−^ strain showed no signs of glucose or ATP depletion when exogenous glucose or oxaloacetate was supplemented in the growth medium. It is noteworthy that the LmCA2^+/−^ strain even under glucose-limiting condition had normal glucose and ATP levels (Table 2). Our data thus provide compelling evidence that LmCA1, but not LmCA2, is involved in gluconeogenesis and ATP production in *L. major*.

**Table 2.** Intracellular glucose and ATP levels of wild type or mutant *L. major* strains grown in low glucose medium in absence or presence of oxaloacetate or glucose.

| Strains/Treatment<sup>a</sup> | Total intracellular glucose (nmol/10<sup>8</sup>cells)<sup>b</sup> | Total intracellular ATP (nmol/10<sup>8</sup>cells)<sup>c</sup> |
| --- | --- | --- |
| WT | 11.71 ± 0.62 | 56.67 ± 5.20 |
| LmCA1<sup>+/-</sup> | 6.80 ± 0.54** | 28.79 ± 2.96** |
| LmCA1<sup>+/-</sup>:CM | 11.49 ± 0.55 | 57.25 ± 2.78 |
| LmCA1<sup>+/-</sup> + 5.6 mM Glu | 12.99 ± 0.65 | 64.37 ± 2.17 |
| LmCA1<sup>+/-</sup> + 5 mM OAA | 12.17 ± 0.58 | 57.46 ± 3.82 |
| LmCA2<sup>+/-</sup> | 12.11 ± 0.62 | 56.06 ± 4.69 |

<sup>a</sup>Wild type (WT) or mutant *L. major* promastigotes were grown in low glucose (0.6 mM) medium for 72 hrs. During this 72 h duration, LmCA1<sup>+/-</sup> strain was also grown in presence of 5 mM oxaloacetate (OAA) or 5.6 mM of glucose (Glu).
<sup>b</sup>Total internal glucose concentration (in nmol) was experimentally determined from 2.5 x 10<sup>8</sup> *L. major* cells grown in low glucose medium.
<sup>c</sup>Total internal ATP concentration (in nmol) was experimentally determined from 8 x 10<sup>6</sup> *L. major* cells grown in low glucose medium.
± indicates SD of values from triplicate experiments.
\*indicates significant difference (\*\*p<0.01) with respect to wild type strain.

**Fig. 2.**
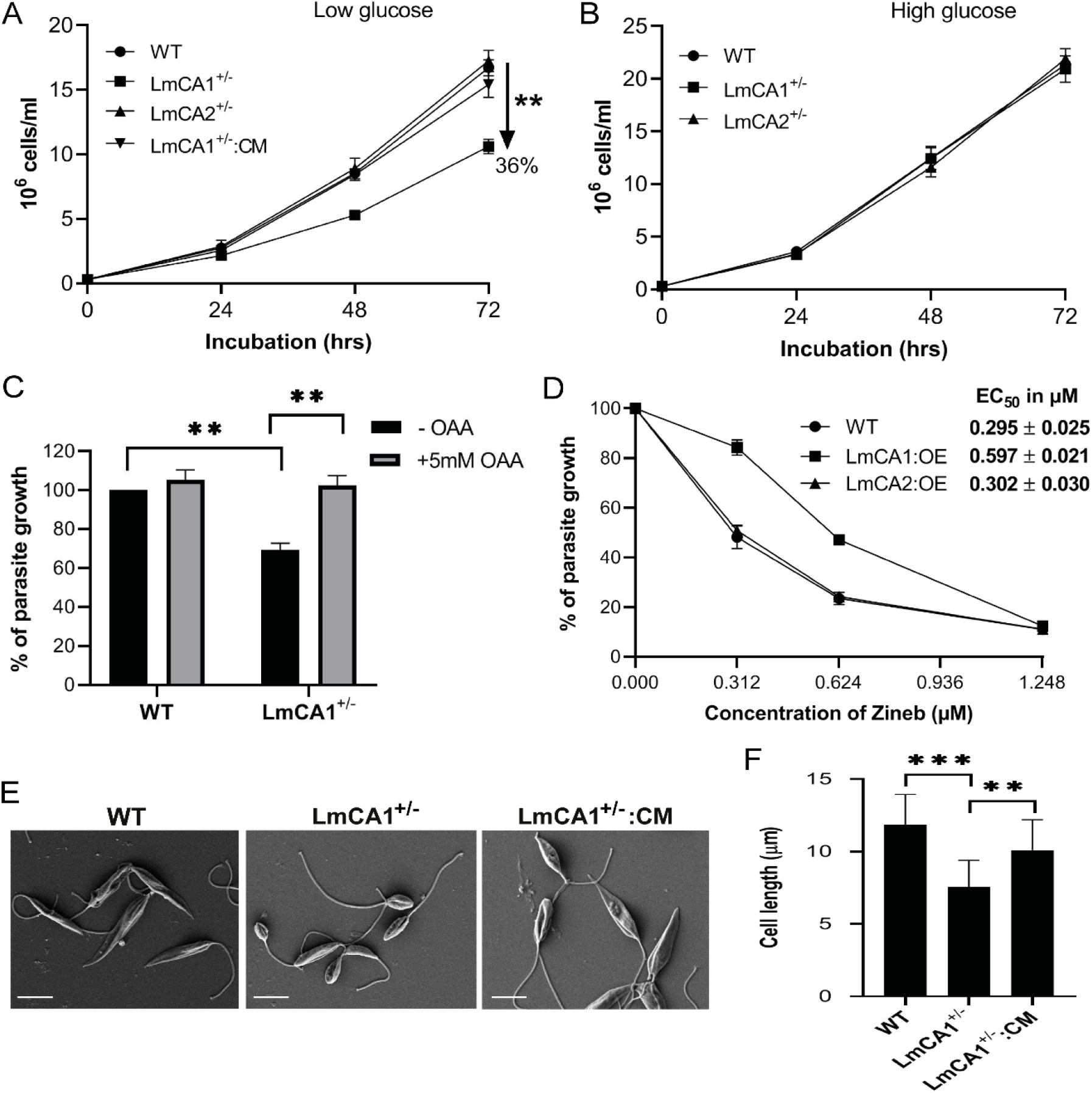
Dissecting the gluconeogenic role LmCA1 and LmCA2. (A) Wild type (WT; circle), LmCA1^+/−^ (square), LmCA2^+/−^ (triangle), and LmCA1^+/−^:CM (inverted triangle) strains were grown in low (0.6 mM) glucose medium, and their growth was monitored every 24 hrs until 72 hrs. A 36% reduction in LmCA1^+/−^cell growth, in comparison to WT, is indicated by a black arrow. Each data point represents the mean result± SD from 3 independent experiments. Asterisk indicates significant difference with respect to WT. **P<0.01 (Student’s t-test). (B) Wild type (WT; circle), LmCA1^+/−^ (square), and LmCA2^+/−^ (triangle) strains were grown in high (6.2 mM) glucose medium, and their growth was monitored every 24 hrs until 72 hrs. Each data point represents the mean result ± SD from 3 independent experiments. (C) WT and LmCA1^+/−^strains were grown in low (0.6 mM) glucose medium, in absence (black bars) or presence (grey bars) of 5 mM oxaloacetate (OAA), and the cell count was performed after 72 hrs of growth. Each data point represents the mean result± SD from 3 independent experiments. Asterisks indicate significant difference with respect to WT (**P≤0.01; Student’s t-test) or LmCA1^+/−^ (**P≤0.01; paired t-test) strain grown in absence of OAA. (D) Wild type (WT; circle), LmCA1-overexpressing (LmCA1:OE; square) or LmCA2-overexpressing (LmCA2:OE; triangle) promastigotes were grown in low (0.6 mM) glucose medium in absence or presence of the indicated concentrations of zineb, and growth of the cells were measured by haemocytometer-based cell counting after 72 hrs of growth. Cell growth of untreated samples was considered as 100%. The EC_50_ values (in μM) of zineb for each *L. major* strain are given in the index. Error bars represent mean ± SD of values from 3 independent experiments. (E) SEM images (6000×) of wild type (WT), LmCA1^+/−^ mutant or LmCA1^+/−^:CM complementation *L. major* promastigotes grown in low (0.6 mM) glucose medium for 72 hrs. Scale bars: 5 μm. (F) Representative bar graph comparing cell length (in μm) of the wild type and mutant strains. Error bars represent average cell length± SD of values. Asterisks indicate significant difference with respect to WT (***P<0.001) or LmCA1^+/−^ (**P<0.01) strain (Student’s t-test).

### Functional cooperation between LmCA1 and LmME in promoting gluconeogenesis and in sustaining parasite growth under glucose-limiting condition

Although our results provided unambiguous evidence to support the role of LmCA1 in gluconeogenesis, it remains unclear how it participates in the process. It is well established that CAs facilitate various metabolic reactions by providing the crucial HCO_3_^−^ to different carboxylating enzymes [38]. However, absence of the PC gene in *Leishmania* genome raises question about the possible metabolic partner of LmCA1 [29,30]. How oxaloacetate could be synthesized bypassing the pyruvate carboxylase reaction was also a mystery. We were intrigued to come across few prior studies where pyruvate carboxylating activity of ME was reported in heart, skeletal muscle, neuronal cells as well as in plant [33,35,36,39]. This led us to hypothesize that LmCA1 might facilitate bicarbonate-dependent pyruvate carboxylation by a leishmanial ME (Fig. 3A). This notion was strengthened by the fact that a functional ME from *L. major* has recently been identified by Giordana *et. al*. Although kinetic and structural properties of this enzyme (henceforth referred to as LmME) were reported, its physiological function is still unknown [37]. Thus, in order to study the physiological function of this enzyme and its possible role in gluconeogenesis we first looked for a potent pharmacological inhibitor of LmME. A high throughput screening by Ranzani *et. al.* led to identification of several inhibitors against two ME isoforms of *Trypanosoma cruzi* [40]. From this large list of inhibitors, we procured three compounds (ATR4-003, ATR6-001, and ATR7-010) based on their high efficiency in inhibiting the *T. cruzi* MEs and potent trypanocidal activity (Fig. S2) [40]. To test whether these compounds can also act as LmME inhibitors, we first cloned LmME cDNA in pET28a vector, expressed the protein in *E. coli* BL21(DE3) and eventually purified it to homogeneity (Fig. S3, 3B). The purified LmME catalyzed NADP^+^-dependent decarboxylation of malate to pyruvate as well as NADPH-dependent carboxylation of pyruvate to malate as reflected by specific activity data (Fig. 3C). Purification of LmME in active form thus allowed us to test efficacies of the potential inhibitors. We found that although ATR4-003 and ATR6-001 did not inhibit LmME activity up to a concentration of 20μM (data not shown), ATR7-010 could inhibit malate decarboxylating and pyruvate carboxylating activity of LmME with IC_50_ values of 1.595μM and 1.556μM, respectively (Fig. 3D). Encouraged by this finding, we next checked if ATR7-010 treatment could affect *L. major* growth. Data presented in Fig. 3E shows that ATR7-010 treatment caused a dose dependent inhibition of parasite growth in glucose-deprived condition with an EC_50_ value of 13.7μM. To check if this stunted growth is indeed due to LmME inhibition, we generated a *L. major* strain overexpressing LmME (LmME:OE) (Fig. S1). As expected, LmME:OE strain exhibited significantly less susceptibility to ATR7-010-mediated growth inhibition under glucose-deprived condition (with EC_50_ value of 24.2μM as compared to 13.7μM for the wild type). Interestingly, ATR7-010-mediated growth inhibition of the wild type *L. major* could be prevented to a significant extent by exogenous addition of glucose or oxaloacetate (EC_50_ values increased by > 2.5 folds in both the cases). These results gave us the first indication that similar to LmCA1, LmME might also be playing an important role in gluconeogenesis. To investigate whether there is any functional cooperation between LmME and LmCA1 we checked for ATR7-010 susceptibility of the LmCA1 overexpressing *L. major*. Strikingly, LmCA1:OE strain showed almost two folds decrease in susceptibility to the LmME inhibitor, ATR7-010, under glucose-deprived condition (Fig. 3E). To crosscheck the functional cooperation between LmME and LmCA1, we performed the reverse experiment whereby we determined susceptibility of the LmME overexpressing strain to LmCA inhibitor, zineb. We found that indeed LmME overexpression provided significant protection against zineb-mediated growth inhibition under glucose limiting condition (EC_50_ values were 0.295μM and 0.722μM for wild type and LmME:OE strains, respectively). Taken together, these results indicate that LmCA1 functionally cooperates with LmME to promote gluconeogenesis and *Leishmania* cell growth under glucose limiting environment. Treatment with ATR7-010 not only affected *Leishmania* growth, but also induced significant morphological deformities resulting in dose dependent shortening of cell length (Fig. 3G, H). Such crippled morphology is an indicator of metabolic stress, which became more obvious when we measured intracellular glucose and ATP levels in the ATR7-010 treated parasites. Data presented in Table 3 shows that there was >30% drop in intracellular glucose and ATP levels in the ATR7-010 treated *L. major* cells as compared to their untreated counterpart. Exogenous supplementation with glucose or oxaloacetate completely protected against depletion of these key metabolites thereby providing strong evidence supporting the involvement of LmME in gluconeogenesis. It is worth noting that intracellular glucose and ATP contents were significantly more in the LmME:OE strain compared to that in the wild type *L. major*. This data suggests that efficiency of gluconeogenesis in the parasite can be controlled by tweaking LmME expression.

**Table 3.** Intracellular glucose and ATP levels of untreated, ATR7-010-treated wild type, or mutant *L. major* strains grown in low glucose medium in absence or presence of glucose or oxaloacetate.

| Strains/Treatment<sup>a</sup> | Total intracellular glucose (nmol/10<sup>8</sup>cells)<sup>b</sup> | Total intracellular ATP (nmol/10<sup>8</sup>cells)<sup>c</sup> |
| --- | --- | --- |
| Untreated WT | 11.71 ± 0.62 | 56.67 ± 5.20 |
| WT + 25µM ATR7-010 | 7.17 ± 0.76** | 38.42 ± 3.79** |
| WT + 25µM ATR7-010 + 5.6 mM Glu | 12.33 ± 0.63 | 63.25 ± 4.56 |
| WT + 25µM ATR7-010 + 5 mM OAA | 13.40 ± 0.51 | 62.08 ± 4.02 |
| LmME:OE | 21.78 ± 0.53 | 98.75 ± 5.25 |

<sup>a</sup>Wild type (WT) or LmME expressing (LmME:OE) *L. major* promastigotes were grown in absence or presence of 25 µM ATR7-010 in low glucose (0.6 mM) medium for 72 h. ATR7-010-treated WT cells were either unsupplemented or supplemented with 5.6 mM glucose (Glu) or 5 mM oxaloacetate (OAA) during the 72 h duration.
<sup>b</sup>Total internal glucose concentration (in nmol) was experimentally determined from 2.5 x 10<sup>8</sup> *L. major* cells grown in low glucose medium.
<sup>c</sup>Total internal ATP concentration (in nmol) was experimentally determined from 8 x 10<sup>6</sup> *L. major* cells grown in low glucose medium.
± indicates SD of values from triplicate experiments.
\*indicates significant difference (\*\*p<0.01) with respect to untreated wild type strain.

**Fig. 3.**
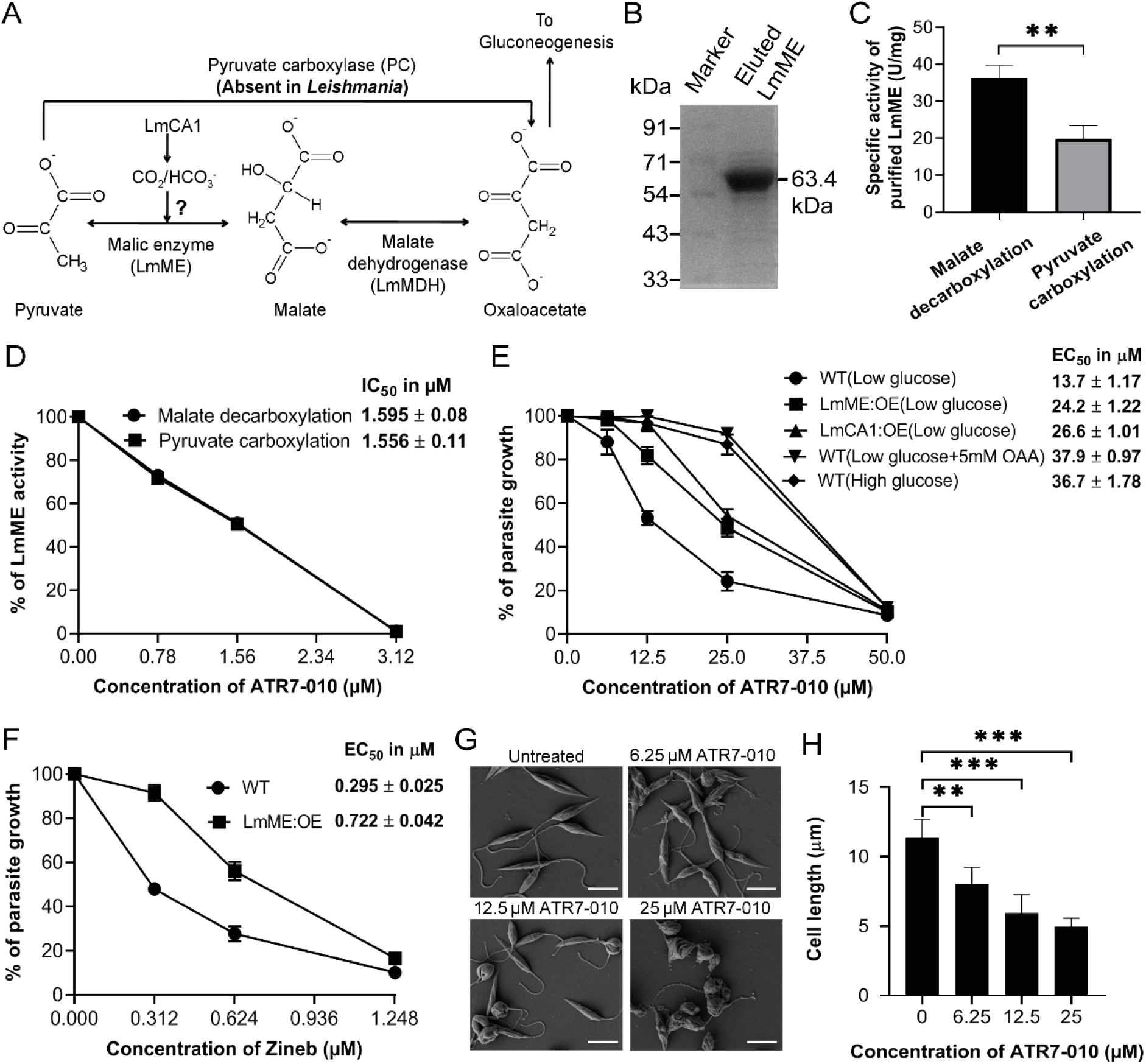
Combined action of LmME and LmCA1 in promoting gluconeogenesis in *L. major* (A) Schematic representation of a plausible PC-independent bypass gluconeogenic pathway in *Leishmania* employing LmCA1, LmME and LmMDH. (B) 10μg of purified 6xHis-tagged LmME protein (63.4 kDa) was loaded on 10% SDS-PAGE, and detected by coomassie blue staining. (C) Bar graph showing malate decarboxylation (black bar) or pyruvate carboxylation (grey bar) activity (in U/mg) in purified LmME measured spectrophotometrically. 5μg of purified LmME was used per assay reaction. Error bars represent mean ± SD of values from 3 independent experiments. Asterisk indicates significant difference between malate decarboxylation and pyruvate carboxylation activity in purified enzyme. **P<0.01(Paired t-test). (D) Malate decarboxylation (circle) or pyruvate carboxylation (square) activity was measured in purified LmME in absence or presence of indicated concentrations of ATR7-010. Enzyme activity in absence of ATR7-010 (0 μM) was considered as 100%. Error bars represent mean ± SD of values from 3 independent experiments. The IC_50_ values (in μM) for ATR7-010 for malate decarboxylation or pyruvate carboxylation activity are given in the index. (E) Wild type (WT; circle), LmME-overexpressing (LmME:OE; square) or LmCA1-overexpressing (LmCA1:OE; triangle) *L. major* strains was grown in low (0.6 mM) glucose medium in absence or presence of increasing concentrations of ATR7-010, and cell number was counted microscopically at 72 hrs. During this period, ATR7-010-treated WT cells were also grown in presence of 5 mM oxaloacetate (OAA; inverted triangle) or 5.6 mM glucose (diamond). Cell number of untreated (0 μM ATR7-010) cells was considered as 100% for each experimental set. Error bars represent mean ± SD of values from 3 independent experiments. Respective EC_50_ values (in μM) are given in the index. (F) Wild type (WT; circle) or LmME-overexpressing (LmME:OE; square) strain was grown in low (0.6 mM) glucose medium in absence or presence of increasing concentrations of zineb, and cell number was counted microscopically at 72 hrs. Cell number of untreated (0 μM zineb) cells was considered as 100% for each experimental set. Error bars represent mean ± SD of values from 3 independent experiments. Respective EC_50_ values (in μM) are given in the index. (G) SEM images (6000×) of wild type *L. major* promastigotes grown in low (0.6 mM) glucose medium in absence (untreated) or presence of indicated concentrations of ATR7-010. (H) Representative bar graph comparing average cell length (in μm) of promastigotes grown in absence or presence of indicated concentrations of ATR7-010. Error bars represent average cell lengths ±SD of values from at least 50 independent measurements. Asterisks indicate significant difference with respect to untreated cells. ***P<0.001, **P<0.01(Paired t-test).

### LmME expression and activity is regulated by glucose

Following the lead from our finding that genetic overexpression LmME could result in increased production of glucose and ATP, we next explored whether LmME expression can be regulated under physiological context. Glucose has been widely reported to be a physiological regulator of gene expression in many cell types in which glucose signalling pathways serves as a fundamental mechanism to optimize different metabolic activities [41–43]. In fact, glucose starvation was shown to upregulate the gluconeogenic enzyme PEPCK in *L. donovani* [20]. This led us to check if LmME expression is regulated by glucose. From our RTqPCR results it is clear that the LmME transcript levels remained unaltered irrespective of the amount of glucose present in the growth medium (Fig. 4A). To analyze LmME expression at the protein level we first generated a rabbit polyclonal antibody againstthe protein (Fig. S4A). We used this antibody to analyze LmME protein levels in *L. major* promastigotes growing in glucose-limiting (low glucose) or glucose-supplemented (high glucose) medium by western blot. In contrast to our RTqPCR data, we observed that LmME protein level was significantly upregulated (more than two folds) in the parasites growing under low glucose condition than those in high glucose (Fig. 4B). The status of LmME was independently verified by immunofluorescence staining, the result of which are in agreement with the western blot data (Fig. 4C). It is worth noting that varying glucose concentration in growth medium did not alter LmCA1 expression either at mRNA or at protein level as determined by RTqPCR and western blot analysis using anti-LmCA1 antibody (Fig. S4B, S5). We next compared LmME activities in whole cell lysate of parasites growing in glucose-limiting or glucose-supplemented medium. We observed that although malate decarboxylating activity remained unaltered, there was ~ 20% increase in pyruvate carboxylating activity of LmME in *Leishmania* cells growing in glucose-limiting condition as opposed to those having access to exogenously added glucose. An even more striking observation was that while malate decarboxylating activity was ~1.5 folds more than the pyruvate carboxylating activity in the purified LmME, the enzyme functioned quite differently when its specific activity was measured in the whole cell lysate. We found that pyruvate carboxylating activity of LmME in *L. major* whole cell lysate is ~5 folds more than the malate decarboxylating activity (Fig. 3C, 4F). Taken together our data suggest that intracellular LmME, especially under low glucose condition, promotes gluconeogenesis by increased protein expression and by selectively augmenting its pyruvate carboxylase activity.

**Fig. 4.**
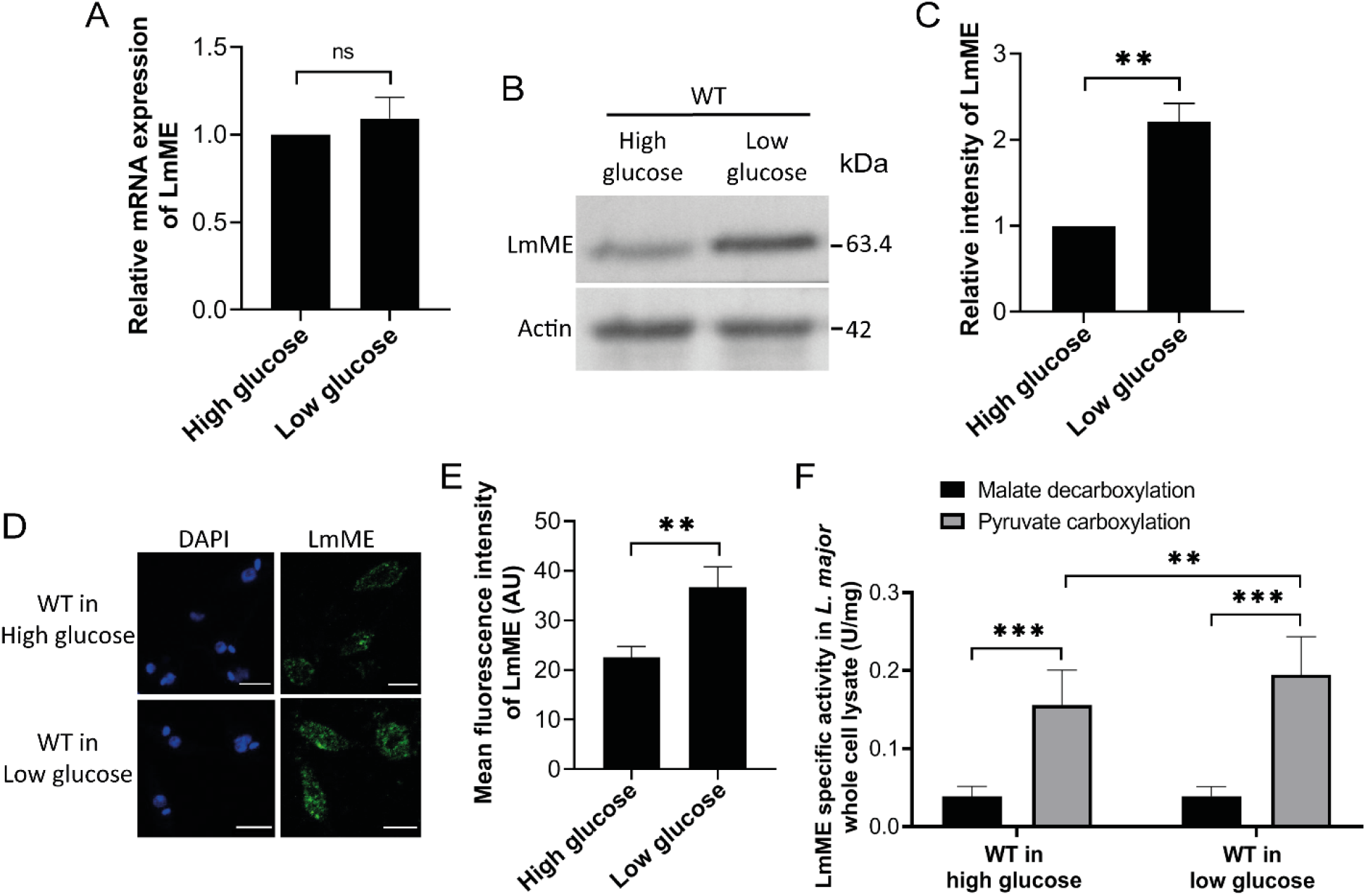
Glucose-mediated regulation of LmME expression and activity. (A) Bar graph showing LmME transcript level in wild type *L. major* promastigotes, grown in high (6.2 mM) or low (0.6 mM) glucose medium for 72 hrs as determined by RTqPCR using rRNA45 as endogenous control gene and cells grown in high glucose condition as reference sample. ‘ns’ indicates insignificant difference, P>0.05 (Paired t-test). (B) Protein level of LmME (63.4 kDa) in whole cell lysates of wild type (WT) *L. major* promastigotes, grown in high (6.2 mM) or low (0.6 mM) glucose medium for 72 hrs, was checked by western blotting using antibody against LmME. Expression of β-actin (42 kDa), detected byanti-β-actin antibody was considered as the loading control. (C) Bar graph comparing relative LmME band intensity of wild type (WT) strain grown in high or low glucose medium from triplicate experiments. Error bars represent mean ± SD of values from 3 independent experiments. Asterisk indicates significant difference with respect to WT cells grown in high glucose medium. **P<0.01 (Paired t-test). (D) Wild type (WT) *L. major* promastigotes, grown in high (6.2 mM) or low (0.6 mM) glucose medium for 48 hrs, were immunostained with an antibody against LmME (green), and visualised with a Zeiss LSM 710 confocal microscope using appropriate filter sets. DAPI (blue) was used to stain the nucleus. Scale bars: 5 μm. (E) Representative bar graph comparing mean LmME fluorescence intensity (in arbitrary units; AU) of wild type (WT) strain grown in high or low glucose medium. At least 50 *L. major* cells were analyzed per experimental condition. Error bars represent mean ± SD of values from 3 independent experiments. Asterisk indicates significant difference in mean fluorescence intensity between WT cells grown in high and low glucose condition. **P<0.01 (Paired t-test). (F) Malate decarboxylation (black bars) or pyruvate carboxylation (grey bars) activity (in U/mg) was spectrophotometrically measured in whole cell lysate of wild type *L. major* promastigotes grown in high (6.2 mM) or low (0.6 mM) glucose medium for 72 hrs. 100 μg of whole cell lysate was used per assay reaction. Error bars represent mean ± SD of values from 3 independent experiments. Asterisk indicates significant difference between malate decarboxylation and pyruvate carboxylation activity, or between pyruvate carboxylation activity in whole cell lysates of wild type *L. major* promastigotes grown in high or low glucose medium.**P<0.01, ***P<0.001 (Paired t-test).

### LmME is localized in the mitochondria

As it is evident from our data that there is a functional cooperation between LmCA1 and LmME in triggering gluconeogenesis, precise knowledge about subcellular localization of these two proteins is of utmost importance. We have previously reported LmCA1 to be a cytosolic enzyme, however, localization of LmME is yet to be determined [28]. To predict subcellular localization of LmME we analyzed its sequence using various bioinformatics tools. The reports predicted LmME to be a mitochondrial protein devoid of any transmembrane domain (Table S1). To experimentally determine its subcellular localization, we first developed *L. major* stable transfectants expressing C-terminal GFP-tagged LmME and stained these cells with mitochondria specific marker, Mito Tracker red (Fig. S6). Extensive co-localization of GFP puncta with the Mito Tracker stained vesicles was observed, suggesting mitochondrial localization of LmME (Fig. 5A).While this initial result was promising, we developed an anti-LmME antibody in the meantime and decided to use it to validate localization of the endogenous LmME in wild type *L. major* by biochemical methods. For this, we lysed the cells with digitonin and separated the whole cell lysate into mitochondrial and cytoplasmic fractions. Western blot of the cell fractions with anti-LmME antibody revealed that LmME is exclusively localized in the mitochondrial fraction. Authenticities of the cell fractionations were confirmed by western blots with antibodies against previously reported mitochondrial (LmAPX) and cytosolic (LmCA1) proteins of *L. major* (Fig. 5B, S4B) [28,44]. Although our data provided unambiguous evidence in support of mitochondrial localization of LmME, its submitochondrial localization (matrix/membrane) could not be ascertained experimentally due to unavailability of appropriate *Leishmania* specific antibody markers. However, since LmME1 lacks any transmembrane domain, it is likely to be localized in the mitochondrial matrix (Table S1).

**Fig. 5.**
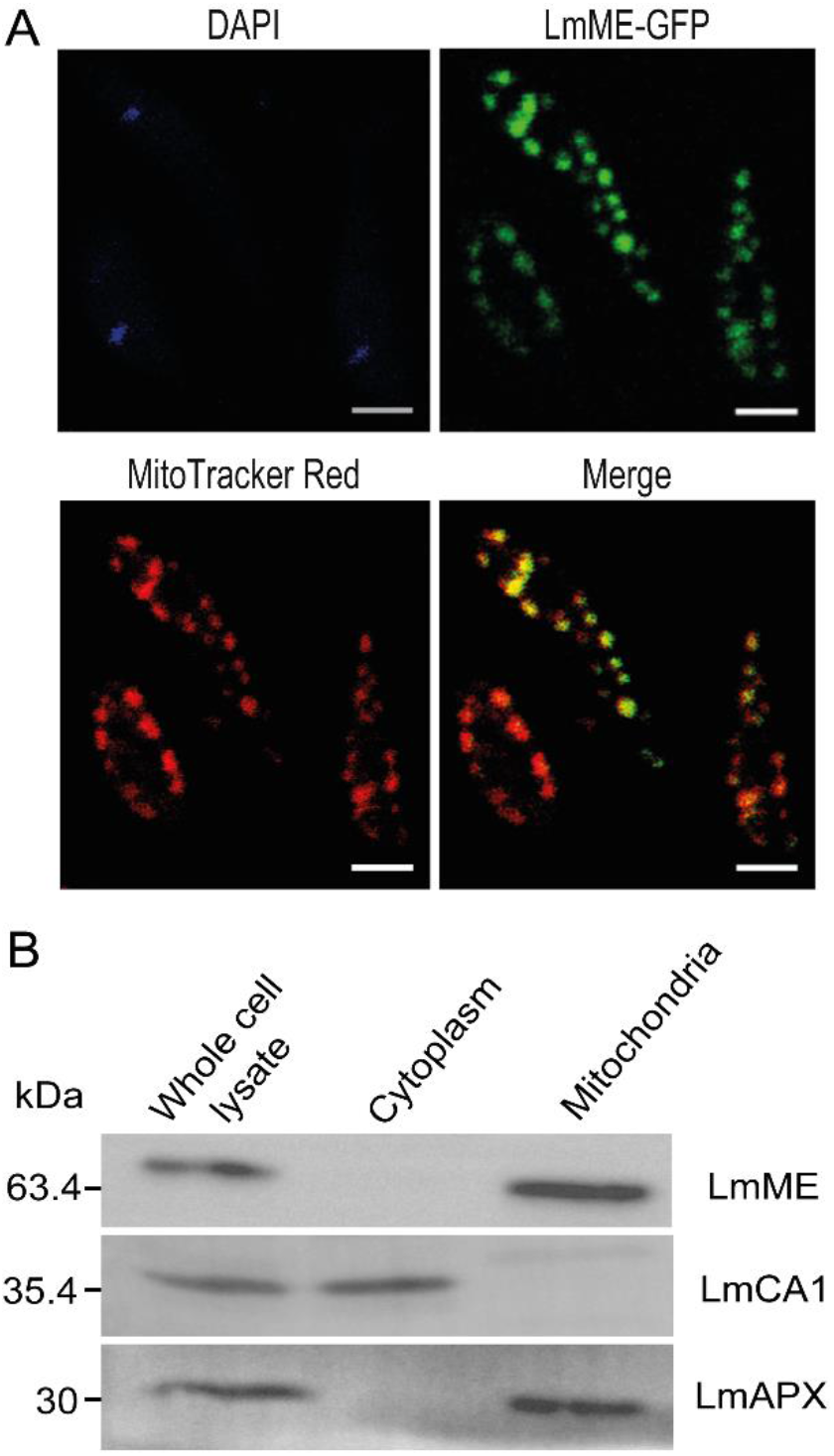
Subcellular localization of LmME. (A) *L. major* cells stably expressing LmME as a C-terminal GFP-tagged protein (LmME–GFP) were stained with MitoTrackerRed CMXRos (red).The pattern of LmME and MitoTracker Red colocalisation (merge) was visualised with a Zeiss LSM 710confocal microscope using appropriate filter sets. DAPI (blue) was used to stain the nucleus. These are representative images from multiple experiments. Scale bars: 2 μm. (B) Wild type *L. major* cells were subjected to fractionation into cytoplasmic and mitochondrial fractions. Distribution of LmME (63.4 kDa) in the cell fractions was determined by western blotting with an antibody against LmME. Authenticity of the cell fractions was verified by western blotting using antibodies LmCA1 (as cytosolic marker, 35.4 kDa) and LmAPX (as mitochondrial marker, 30 kDa). Shown are representative blots from at least 3 independent experiments.

### LmCA1 and LmME are both crucial for survival of *L. major* within host macrophages

After having proven that LmCA1 cooperates with LmME in sustaining *in vitro* growth of the parasites under glucose limiting condition, we wanted to check the importance of these two enzymes for intracellular propagation of *L. major*. For this, wild type *L. major* were first grown under glucose limiting condition, following which we infected J774A.1 macrophages with these parasites in absence or presence of 0.625μM zineb or 25μMATR7-010. It was found that in presence of LmCA1 inhibitor (zineb) and LmME inhibitor (ATR7-010) there was 43% and 35% drop in the intracellular parasite burden, respectively as compared to untreated control (Table 4). Based on our previous report and the data presented in Fig. S7, it is worth pointing out that the concentration of inhibitors used in this experiment (0.625μM for zineb and 25μM for ATR7-010) do not have any effect on macrophage growth [27]. Next, J774A.1 macrophages were infected with the corresponding overexpressing strains of *L. major* (LmCA1:OE and LmME:OE). In contrast to the results obtained with the enzyme inhibitors, we observed ~55% spike in intracellular parasite burden for both the strains as compared to their wild type counterpart (Table 4). Taken together, our data confirmed that LmCA1 as well as LmME are indeed very critical for intracellular propagation of *L. major*. Interestingly, in all these experimental conditions there was not much difference in the percentage of infected macrophages, which varied from 86 – 96%, suggesting that LmCA1 and LmME possibly do not play any major role in infectivity of the parasite (Table 4).

**Table 4.** Intracellular parasite burden of glucose-starved *L. major* cells.

| Strains/Treatment<sup>a</sup> | Amastigotes/100 macrophages<sup>b</sup> | Percentage of infected macrophages<sup>c</sup> |
| --- | --- | --- |
| Untreated WT | 341.33 ± 24.84 | 91.56 ± 1.32 |
| WT + 0.625 µM Zineb | 196.33 ± 16.01** | 86.24 ± 1.19 |
| WT + 25 µM ATR7-010 | 224 ± 23** | 87.69 ± 2.29 |
| LmCA1:OE | 610.66 ± 43.52*** | 96.72 ± 2.07 |
| LmME:OE | 601 ± 41.58*** | 94.80 ± 0.53 |

<sup>a</sup>J774A.1 macrophages were infected with stationary-phase wild type (WT) or overexpressing (LmCA1:OE and LmME:OE) *L. major* strains grown previously in low (0.6 mM) glucose medium for 72 hrs. During the 18 hr-infection period, WT *L. major*-infected J774A.1 macrophages were treated with 0.625 µM zineb or 25 µM ATR7-010.
<sup>b</sup>Number of amastigotes per 100 macrophages (parasite load) for each strain/treatment was quantified by counting the total number of DAPI-stained nuclei of macrophages and amastigotes in a field. For each condition, atleast 100 macrophages (and corresponding number of amastigotes) were analysed.
<sup>c</sup>Percentage of macrophages infected by each *L. major* strain/treatment was determined by counting total number of DAPI-stained nuclei of uninfected and infected macrophages in a field. For each condition, atleast 100 macrophages were analysed.
± indicates SD of values from triplicate experiments.
\*indicates significant difference (\*\*P<0.01, \*\*\*P<0.001) with respect to untreated WT.

## Discussion

Ability to synthesize glucose from non-carbohydrate precursors via gluconeogenesis is the mainstay for survival of several organisms under sugar limiting condition [45]. This metabolic pathway is particularly important for the *Leishmania* parasites, which grows within the amino acid rich phagolysosomal compartment [12,17,46]. Although some of the gluconeogenic enzymes of *Leishmania* were earlier identified through elegant studies, very little is known about the details of the entire pathway [18,19]. Absence of the PC encoding gene in *Leishmania* genome was especially perplexing and hence the mechanism of pyruvate entry into the gluconeogenic circuit remained elusive till date [29]. Our work revealed that LmME, by virtue of its unconventional pyruvate carboxylating activity, drives gluconeogenesis in *L. major* with help of the CO_2_ concentrating enzyme, LmCA1. This unique functional partnership between cytosolic LmCA1 and mitochondrial LmME was found to be critical for growth of *Leishmania* promastigotes when glucose availability was restricted. Both of these enzymes also played determining roles in establishing intracellular *Leishmania* infection within host macrophages. Thus, apart from providing new mechanistic insights into the gluconeogenic pathway of *Leishmania*, our results highlights LmCA1 and LmME as prospective drug targets, worthy of further exploration.

The first step of gluconeogenesis in mammals involves conversion of pyruvate to oxaloacetate by the mitochondrial enzyme PC. This is a critical entry point through which several gluconeogenic amino acids are funnelled into the *de novo* glucose synthesis pathway [22,45]. PC is a biotin-dependent carboxylase that uses HCO_3_^−^ as the donor of the carboxyl group [22]. Mitochondrial CAV is known to facilitate the PC-catalyzed carboxylation reaction by providing this crucial HCO_3_^−^ [24,26]. Although CAV is a well-established player of mammalian gluconeogenesis, there is no information regarding gluconeogenic capability of CAs in lower vertebrates, invertebrates or in microorganisms [38]. In this context, our data showing the role of LmCA1 in supporting gluconeogenesis in *L. major* is possibly the first evidence of gluconeogenic activity of a CA in lower eukaryotes. This exciting result was, however, difficult to comprehend because *Leishmania* genome does not encode a *bona fide* PC gene [29,30]. How LmCA1 participates in the gluconeogenesis process in absence of its metabolic partner PC was an intriguing question for us.

It was earlier proposed that in absence of PC, the parasite may utilize the enzyme pyruvate phosphate dikinase (PPDK) to directly synthesize phosphoenolpyruvate (PEP) from pyruvate without having to go through the oxaloacetate intermediate [19]. But experiments with Δ*PPDK* mutant *Leishmania* confirmed that PPDK do not play any role in gluconeogenesis in *Leishmania* promastigotes. Although PPDK was shown to be responsible for pyruvate entry into the gluconeogenic pathway in axenic amastigotes, functional importance of this enzyme is yet to be established in intracellular amastigotes [19]. Thus, it is evident that the available information on PPDK function in *Leishmania* fails to provide a comprehensive understanding of the PC-independent mechanism of pyruvate entry into the gluconeogenic pathway. Neither does it clarify the exact role of LmCA1 in this process. These ambiguities were finally removed with the identification of LmME as an important player of the gluconeogenesis in *L. major*. Our data suggests that pyruvate carboxylating activity of LmME provides an alternative mode of pyruvate entry into the gluconeogenic pathway via pyruvate – malate – oxaloacetate route. This is supported by the observation that pyruvate-carboxylation was the dominant activity of LmME in *L. major* whole cell lysate. This is in stark contrast to the purified enzyme, in which the malate-decarboxylating activity was found to be dominant. This interesting data suggests that some unknown factors in *Leishmania* cells may act as LmME regulator in promoting its pyruvate carboxylating activity *in vivo*.

ME is a ubiquitous enzyme that catalyzes reversible decarboxylation of malate in presence NADP to produce pyruvate, CO_2_ and NADPH [47]. Although the enzyme exhibits both carboxylating and decarboxylating activity when assayed *in vitro*, malate-decarboxylation was shown to be responsible for most of the reported physiological function of ME. The reducing equivalent (NADPH), generated as a byproduct of this decarboxylation reaction, was shown to promote fatty acid biosynthesis and maintain redox balance in various organisms [48–52]. Relatively much less is known regarding the physiological role of the pyruvate-carboxylating activity of ME. Hassel B *et. al.* reported that ME-catalyzed pyruvate carboxylation in rat neurons results in synthesis of TCA cycle intermediates that in turn promotes production of the neurotransmitter glutamate [33]. Apart from this, pyruvate-carboxylating activity of ME was shown to play a role in anaplerosis in hypertrophied heart as well as in plant (*Arabidopsis thaliana*) whereby malate produced in the cytosol is transported to mitochondria for fuelling the TCA cycle [35,36]. Our finding showing participation of LmME in gluconeogenesis in *Leishmania* through its pyruvate-carboxylating activity thus uncovers a novel physiological function of ME.

Functional partnership between LmME and LmCA1 is another interesting revelation of this study. That the adverse effect of LmCA1 inhibition on *L. major* cell growth under glucose limiting condition could be prevented to a significant extent by overexpression of LmME (and vice versa) is a testimony of the fact that these two enzymes indeed cooperates with each other. CAs are known to play a crucial role in metabolism by providing the CO_2_/bicarbonate to various carbon fixing enzymes (e.g. PC, carbamoyl phosphate synthetase, RuBisCO) that incorporates CO_2_ to the corresponding substrates [38,53]. Our data suggests that LmCA1 facilitates the carboxylation reaction catalyzed by LmME in a similar way. We have previously reported that the cytosolic LmCA1 is instrumental in HCO_3_^−^ buffering of *Leishmania* cytosol by converting the incoming H^+^ ions into H_2_O and CO_2_ (). CO_2_ being a freely diffusible gas can easily disperse across the mitochondrial membranes having high CO_2_-permeablity and stimulate pyruvate carboxylating activity of LmME in the lumen of the mitochondria to generate malate [54,55]. This appears to be rational mechanism since ME has a substrate preference for CO_2_ as opposed to its counterpart PC, which utilizes HCO_3_^−^ [22,56,57]. Absence of a mitochondrial CA in *L. major* also seems to be a key factor in maintaining a high luminal concentration of CO_2_, which otherwise would have been readily converted to HCO_3_^−^. A similar CO_2_ concentrating mechanism has been reported in the chloroplast of green-alga. It was shown that a CA present in the thylakoid lumen converts bicarbonate to CO_2_, which then diffuses out of the thylakoid double membrane and drives the RuBisCO-catalyzed carboxylation reaction in the stroma [58]. Presence of functionally active malate dehydrogenase (MDH) isoforms in *Leishmania* indicates that the parasite would be able to synthesize oxaloacetate from malate, once the latter is formed by pyruvate carboxylation [59]. Since many of the downstream gluconeogenic enzymes in *Leishmania* and other trypanosomatids (e.g. PEPCK, FBP etc.) are localized in the glycosome, it is likely that the malate formed in the mitochondria is transported to the glycosome before it is converted to oxaloacetate by the catalytically active glycosomal MDH [18,59,60]. The putative malate transporters encoded in the *Leishmania* genome may play an important role in this process by facilitating mitochondria-glycosome malate shuttling [29,30]. A tentative model describing these initial steps of gluconeogenesis in *Leishmania* is outlined in Fig. 6.

**Fig. 6.**
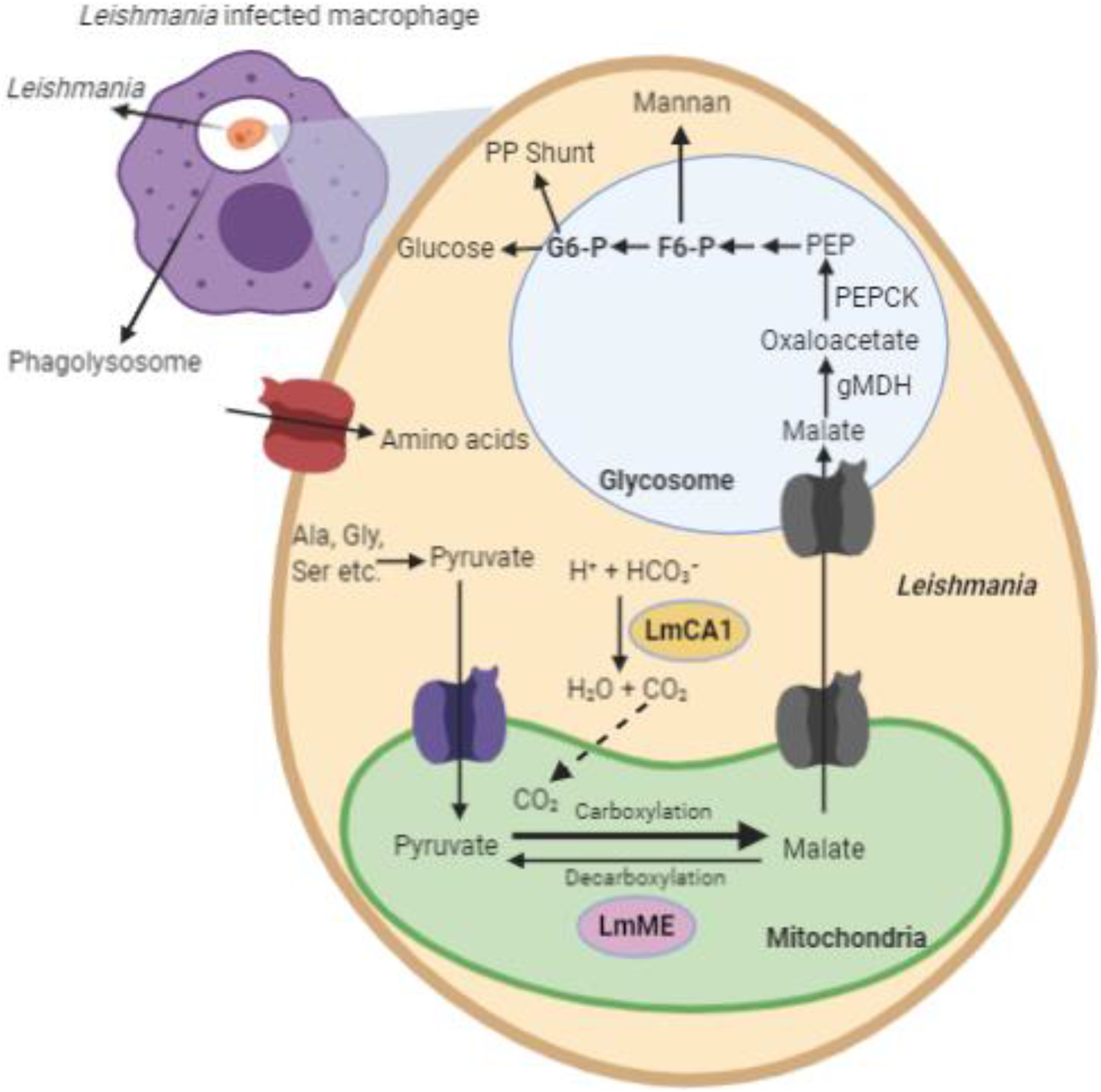
An illustrative model of the initial steps of gluconeogenesis in *Leishmania* highlighting the functional partnership between LmCA1 and LmME. *Leishmania* parasites proliferate in the amino acid-rich phagolysosomal environment of the host macrophages where glucose availability is scarce. Gluconeogenic amino acids, taken up by the parasite from its surroundings, are metabolized to pyruvate in the cytosol. Pyruvate can be transported to the mitochondria through pyruvate carrier protein. Mitochondrial LmME, by virtue of its pyruvate carboxylase activity, can convert pyruvate to malate. The cytosolic LmCA1 can facilitate this carboxylation reaction by producing the crucial CO_2_, which can easily diffuse into the mitochondria. Malate eventually gets transported to the glycosome via putative malate transporters, where it is then converted to oxaloacetate by gMDH. Further downstream pathways of gluconeogenesis shown in the figure are self-explanatory. Abbreviations used: LmCA1; *L. major* carbonic anhydrase 1, LmME; *L. major* malic enzyme, gMDH; glycosomal malate dehydrogenase, PEPCK; Phosphoenolpyruvate carboxykinase, PEP; Phosphoenolpyruvate, F6-P; Fructose 6-phosphate, G6-P; Glucose 6-phosphate, PP shunt; Pentose Phosphate shunt. The image was created using the software, BioRender.com.

In addition to their functional role in *L. major* promastigotes, LmCA1 and LmME were also found to be important for intracellular survival of the amastigotes. *Leishmania* amastigotes resides within the phagolysosomal compartment where glucose availability is limited and amino acids are in abundance [12]. LmCA1-LmME metabolic partnership in promoting *de novo* synthesis of glucose would thus provide a life support for the parasite residing in such a stringent nutritional environment. Apart from their gluconeogenic role, it is also possible that LmCA1 and LmME might have some other physiological function, which can as well contribute to overall fitness of the amastigotes in withstanding the harsh phagolysosomal conditions. In fact, role of LmCA1 in maintaining cytosolic pH homeostasis and acid tolerance of the parasite is already reported by us [28]. Although a non-gluconeogenic-role for LmME has not been reported yet, this possibility cannot be ruled out completely. In this connection it is worth mentioning that while LmME protein level was induced in low glucose concentration, LmCA1 was constitutively expressed. Since glucose responsive expression is a typical characteristic of many gluconeogenic enzymes, it might be speculated that LmME plays a dedicated role in gluconeogenesis whereas function of LmCA1 is more versatile [20,61–63].

To summarize, the metabolic partnership between LmCA1and LmME reported here reveals a novel bypass pathway in gluconeogenesis that allows PC-independent entry of pyruvate into the gluconeogenic circuit in *Leishmania*. Whether this pathway is operational in any other organism remains to be seen. However, it may be noted that both *Trypanosoma brucei* and *Trypanosoma cruzi* lacks the PC gene in their genome and they both express functional CA and ME [29,30,64]. Thus, the CA-ME bypass route may be a characteristic feature of gluconeogenesis for the entire trypanosomatid family. While this will require experimental validation, our current study has clearly established the functional importance of LmCA1 and LmME and uncovered their potential as antileishmanial drug targets.

## Materials and methods

Unless otherwise mentioned, all reagents were purchased from Sigma-Aldrich (St. Louis MO). All primers were bought from Integrated DNA technologies and their sequence details are provided in Table S2.

### Preparation of *Leishmania* culture medium containing high or low glucose concentration

For our study, glucose-free RPMI 1640 (HiMedia) was used, which is rich in gluconeogenic amino acids. It was supplemented with 15% foetal bovine serum (Gibco), 23.5mM HEPES, 0.2mM adenine, 150 μg/ml folic acid, 10 μg/ml hemin, 120 U/ml penicillin, 120 μg/ml streptomycin, and 60 μg/ml gentamicin. This medium was supplemented with or without 5.6 mM glucose, and henceforth been referred to as ‘high glucose’ or ‘low glucose’ medium, respectively. pH was adjusted to 7.2 for both media. Total glucose concentration in high or low glucose culture medium was estimated by glucose oxidase-peroxidase assay, as described later in this section. The glucose concentration for high or low glucose culture medium, was found to be 6.2 mM or 0.6 mM, respectively. Glucose in low glucose medium is contributed by foetal bovine serum.

### Cell culture and cell growth analysis

Wild type *L. major* promastigotes (strain 5ASKH; generously provided by Dr Subrata Adak of IICB, Kolkata, India) were grown at 26°C, as described by us previously [27,28]. J774A.1 (murine macrophage cell line from the National Centre for Cell Science, Pune, India) cells were cultured as described earlier [27,28]. For cell growth analysis, wild type or mutant *L. major* cells were seeded in high (6.2 mM) or low (0.6 mM) glucose medium, and their growth was monitored at different time points till 72 hrs by counting the number of cells in haemocytometer. Wherever mentioned, cells grown in low glucose medium were supplemented with 5 mM oxaloacetate (OAA) or 5.6 mM of glucose (Glu). Selection antibiotics were removed from culture medium during the course of all these experiments. CA inhibitor, zineb [zinc ethylene-bis-dithiocarbamate], or ME inhibitors, ATR4-003 [Pyrimidin-7-one], ATR6-001 [Tetrahydrothieno-isoquinoline] or ATR7-010 [Triazolo-thiadiazole] (ChemBridge Corporation, San Diego, CA), were used for growth inhibition study [40]. The inhibitors were freshly dissolved in dimethyl sulfoxide (DMSO) to prepare 5mM (for zineb) or 100 mM (for ME inhibitors) stock solutions. According to experimental requirements, further dilutions were made in DMSO before addition to the culture medium. *L. major* promastigotes or J774A.1 macrophage were grown in medium containing the inhibitors at desired concentrations for 72 h, following which the cells were analysed using a haemocytometer. Cells incubated with an equivalent concentration of DMSO (0.2%) always acted as untreated controls. The percentage of cell growth was calculated using the formula (Cell number_treated_/Cell number_untreated_×100). The growth of the untreated cells was considered as 100%. Finally, 50% effective concentration (EC_50_) for each inhibitor was calculated from the percentage of cell growth values using OriginPro 8 software. Selection antibiotics were removed from culture medium during the course of all these experiments.

### Transfection

Transfection of DNA into *L. major* cells was performed using electroporation as described by us previously [28]. Briefly, 3.6 × 10^7^ log phase wild type or mutant *L. major* promastigotes were incubated with 10-30 μg of the DNA construct in electroporation buffer (21 mM HEPES,6 mM glucose, 137 mM NaCl and0.7 mM NaH_2_PO_4_; pH 7.4) in a 0.2 cm electroporation cuvette for 10 minutes on ice. Subsequently, electroporation was done in a Bio-Rad Gene Pulsar apparatus using 450 volts and 550 μF capacitance. Transfected cells were selected in appropriate antibiotic-containing medium.

### Generation of *L. major* strain overexpressing LmPEPCK, LmCA1, LmCA2 or LmME

Primers P1/P2, P3/P4, P5/P6 or P19/P20 (listed in Table S2) were used to PCR-amplify the ORF of LmPEPCK, LmCA1, LmCA2 or LmME gene from genomic DNA of wild type *L. major* cells. Amplified LmPEPCK, LmCA1, LmCA2 or LmME gene fragment was cloned into the BamHI/EcoRV sites of pXG-GFP+, SmaI site of pXG-SAT, BamHI site of pXG-PHLEO or BamHI/EcoRV sites of pXG-GFP+ plasmid, respectively, to generate the overexpression (OE) constructs. The clones were subsequently verified by sequencing. 30 μg LmPEPCK:OE, LmCA1:OE, LmCA2:OE or LmME:OE construct was transfected into wild type *L. major* promastigotes by electroporation. Each transfected strain was selected and maintained in 100 μg/mlG418 sulphate, 200 μg/ml nourseothricin (Jena Bioscience), or 8μg/ml phleomycin (Invivogen).

### Generation of *L. major* strain expressing GFP-tagged LmME

Primers P7/P8 (listed in Table S2) were used to PCR-amplify the ORF of LmME gene from genomic DNA of wild type *L. major* promastigotes. Amplified gene segment of LmME was cloned into the BamHI and EcoRV sites of pXG-GFP+ vector to generate the C-terminal GFP-tagged construct. The clone was subsequently verified by sequencing. 30 μg LmME-GFP construct was transfected into wild type *L. major* promastigotes by electroporation. The transfected strain was selected and maintained in 100 μg/mlG418 sulphate.

### Cloning, bacterial expression and purification of LmME

The ORF of LmME gene was PCR-amplified from wildtype *L. major* genomic DNA using the primer set P9/P10. A 1722 bp amplified gene fragment was cloned within EcoRI/HindIII sites of pET28a+ vector to generate the N-terminal 6xHis-tagged construct. The clone was verified by sequencing. For the purpose of protein expression, this construct was transformed into *E. coli* BL21(DE3) cells. Transformed cells were grown overnight in 5ml LB medium containing 50 μg/ml kanamycin at 37°C.Overnight grown culture was inoculated in 250 ml LB medium. When the culture reached an OD_600_ ~0.6, LmME protein expression was induced in presence of 0.5 mM isopropyl β-D-thiogalactoside (IPTG) for 8 hrs at 20°C. Bacterial cells were harvested, resuspended in ice-chilled lysis buffer (50 mM Tris, 100mM NaCl, 10mM imidazole, 1mg/ml lysozyme and 1mM PMSF; pH 8.0) and incubated on ice for 40 min with intermittent vortexing. Cells were lysed using a 10 sec pulse sonicator with 20 sec rest on ice. The cell lysate was subsequently centrifuged at 18000 × g for 30 min at 4°C. The cell free supernatant was loaded on to pre-equilibrated Ni^2+^-nitrilotriacetic resin (Qiagen), and incubated for 1hr at 4°C. The resin was washed with wash buffer (50 mM Tris, 100mM NaCl, 20mM imidazole and 1mM PMSF; pH 8.0), followed by another wash with the same buffer containing 40 mM imidazole. Finally,6xHis-tagged LmME protein bound to Ni^2+^-nitrilotriacetic resin was eluted in wash buffer containing 250 mM imidazole. Eluted LmME was dialyzed thrice in dialysis buffer (50 mM Tris, 100mM NaCland1mM PMSF; pH 8.0). Purity of LmME protein was verified on 10% SDS-PAGE followed by coomassie blue staining.

### LmME and LmCA1 antibody generation

Polyclonal antiserum against LmCA1 or LmME was generated by BioBharati Life Science Pvt. Ltd. (custom antibody generation facility), India. Purified LmME protein (dissolved in sterile PBS, pH 7.4), was used for generating antibody as per company protocol. Briefly, 500 μg of purified LmME was mixed thoroughly with Freund’s Complete Adjuvant (1:1 ratio), and was injected subcutaneously into two adult New Zealand rabbits in equal amounts. After 2 weeks, a booster dose of 150 μg purified protein, mixed with Incomplete Freund’s Adjuvant, was injected into each rabbit. After 5 such booster doses, 10 ml blood was taken from the ear vein of each rabbit, sera were collected and tested by western blotting on purified LmME (0.5 μg/well) and wild type *Leishmania* whole cell lysate (80 μg/well) samples.

The PCR-amplified ORF of LmCA1gene (primers P11/P12 are listed in Table S2) was cloned within EcoRI site of pET28a+ vector, and the verified N-terminal 6xHis-tagged construct was provided to BioBharati Life Science Pvt. Ltd for LmCA1 antiserum generation. LmCA1 protein was induced in BL21(DE3) *E. coli* cells using 0.5 mM IPTG for 4 hrs at 37°C. LmCA1 in the insoluble fraction was used for antigen preparation and administered into adult New Zealand rabbits, as described above. Anti-LmCA1 antiserum was collected and verified by western blotting on LmCA1 (9.25 μg/well) purified to homogeneity from LmCA1-overexpressing *L. major* promastigotes, and wild type *Leishmania* whole cell lysate (120 μg/well) samples.

### LmME activity assay

Malic enzyme activity in purified LmME or *L. major* whole cell lysate was assayed on Hitachi U2900 spectrophotometer using quartz cuvette of 1 cm path length as described previously, with minor modifications [32]. To test malate decarboxylation or pyruvate carboxylation activity, the assay mixture was made up of 1 ml malate buffer (50 mM Tris-Cl; pH 7.5, 10 mM malate, 1 mM MnCl_2_ and 0.15 mM NADP^+^) or pyruvate buffer (50 mM Tris-Cl buffer; pH 5.5, 1 mM MnCl_2_, 0.15 mM NADPH, 50 mM pyruvate and 75 mM NaHCO3), respectively. After incubating in the spectrophotometer at 37°C for 5 min to achieve temperature equilibrium, malate decarboxylation or pyruvate carboxylation reaction was initiated with the addition of 5 μg purified enzyme or 100 μg whole cell lysate. Absorbance was recorded at 340 nm from 0-2min. The average malate decarboxylation or pyruvate carboxylation activity from three different protein preparations or promastigote cultures was expressed in enzyme units (EU)/mg, where 1 unit of enzymatic activity is defined as the amount of enzyme that catalyses production or consumption of 1μmol of NADPH per minute, respectively. Enzyme activity was calculated by considering molar extinction coefficient for NADPH is 6.22 mM^−1^cm^−1^. The total protein concentration of the purified enzyme or whole cell lysate was measured by the method of Lowry *et al* [65]. For inhibition studies, the inhibitors (at desired concentrations) were incubated with purified LmME for 5 mins at room temperature prior to the assay. The 50% enzyme activity inhibitory concentration (IC_50_) for each inhibitor was calculated in triplicate using Origin Pro8.0 program.

### Imaging studies

Morphology of *L. major* promastigotes was determined by Zeiss Supra 55VP scanning electron microscope (SEM) as described by us previously [27,28]. At least 50 cells were analysed for each experimental condition using ImageJ software. During the course of the experiment, selection antibiotics were removed from culture medium.

To determine subcellular localization of LmME in *L. major*, LmME-GFP expressing cells were mounted on poly L-lysine coated coverslips for 1 hr. Attached parasites were then stained with 500 nM MitotrackerRed CMX-Ros (Invitrogen) in the dark for 30 mins [66]. Post-incubation, cells were washed in PBS to remove excess stain, air-dried, and finally embedded in anti-fade mounting medium containing DAPI (VectaShield from Vector Laboratories). Images were acquired with a Zeiss LSM 710 confocal microscope. During the course of the experiment, selection antibiotics were removed from culture medium.

For investigating LmME expression in *L. major*, wild type cells were grown in low (0.6 mM) or high (6.2 mM) glucose medium for 48 hrs. Subsequently, cells were mounted on poly L-lysine coated coverslips, fixed with methanol: acetone (1:1), and permeabilized with 0.1% triton X-100. 0.2% gelatine was used to block non-specific binding. Next, cells were incubated with anti-LmME primary antibody (1:1500) for 1.5 hrs. Cells were washed with PBS and incubated with a secondary goat anti-rabbit Alexa Fluor 488 antibody (1:600; Molecular Probes) for 1.5 hrs in the dark. Post-incubation, cells were washed with PBS and embedded in anti-fade mounting medium containing DAPI. Images were acquired with a Zeiss LSM 710 confocal microscope. Mean fluorescence intensity for different samples was quantified using MacBiophotonics ImageJ software. At least 50 cells were analysed for each set of experiment.

### Subcellular fractionation and western blot analysis

Cytoplasmic and mitochondrial fractions were isolated from wild type *L. major* whole cell lysates as described previously [67,68]. Briefly, 1 × 10^8^ promastigotes were harvested and washed in MES buffer (20mM MOPS, pH 7.0, 250mM sucrose, 3mM EDTA). Cells were resuspended in 0.2 ml MES buffer containing 1 mg/ml digitonin and protease inhibitor cocktail, and incubated at RT for 10 min. The resultant whole cell lysate was centrifuged at10,000 × g for 5 min. The supernatant was collected as the cytoplasmic fraction whereas the pellet was dissolved in MES buffer and used as the mitochondrial fraction.

SDS-PAGE (10%) was performed with the subcellular fractions (sample loaded was equivalent to 5×10^6^cells). LmME was detected with rabbit anti-LmME antisera (1:4000). The authenticity of the cytoplasmic or mitochondrial fraction was verified by western blotting using rabbit anti-*L. major* carbonic anhydrase or -LmCA1 (1:1000) or rabbit anti-*L. major* ascorbate peroxidase or -LmAPX (1:50, a generous gift from Dr Subrata Adak, IICB, India) [44]. After overnight primary antibody incubation at 4°C, blots were probed with anti-rabbit horseradish peroxidase (HRP)-conjugated secondary antibody (1:4000; Thermo Scientific) for 2 hrs. SuperSignal West Pico Chemiluminescent substrate (Thermo Scientific) was used to develop the blots, and chemiluminescent signal was detected in the ChemiDoc imaging system (Syngene).

To investigate relative expression of LmME (63.4 kDa) or LmCA1 (35.4 kDa) in wild type *L. major*, 1 × 10^8^cells were grown in high (6.2 mM) or low (0.6 mM) glucose medium, harvested, resuspended in 200 μl 1X PBS (containing 1 mM PMSF) and lysed by sonication. SDS-PAGE (10%) was performed with samples obtained from 5×10^6^ cells. LmME or LmCA1 was detected with rabbit anti-LmME or -LmCA1 antiserum, as described earlier. Expression of β-actin, detected by rabbit anti-*L. donovani* β-actin antibody (1:4000; a generous gift by Dr Amogh Sahasrabuddhe, CSIR-CDRI) and anti-rabbit HRP-conjugated secondary antibody (1:4000),was considered as the endogenous control [69]. Densitometry value of LmME protein bands was quantified using MacBiophotonics ImageJ software. To check LmME purified in bacterial expression system, 10 μg of purified protein was loaded on 10% SDS-PAGE and detected by western blotting using primary anti-His antibody (1:2000; Bio Bharati Life Science Pvt. Ltd.), followed by anti-rabbit HRP-conjugated secondary antibody (1:4000).

### Measurement of glucose concentration

Intracellular glucose in *L. major* promastigotes was measured by an end-point colorimetric assay, involving the sequential catalytic actions of glucose oxidase (GOD) and peroxidase (POD) enzymes, as described previously with minor modifications [70,71]. 2.5×10^8^ *L. major* cells, grown in low (0.6 mM) glucose medium, were harvested, resuspended in PBS and lysed by sonication. Subsequently, glucose assay solution (98% GOD-POD reagent and 2% o-dianisidine) was added to whole cell lysate and the entire mixture was incubated for 30 mins at 37°C. After incubation, the reaction was stopped by adding12 N H_2_SO_4_, which also allowed formation of a stable coloured product. Finally, the absorbance of samples was measured against the reagent blank at 540 nm. The glucose concentration for each sample was obtained from the corresponding absorbance value using a calibration curve. The calibration curve was generated by recording absorbance as a function of glucose concentration by checking the absorbance of increasing concentrations of glucose (0.625-20μM). Selection antibiotics were removed from culture medium during the course of the experiment. Glucose concentration in the *L. major* culture medium was also measured using this method.

### Determination of intracellular ATP content

Intracellular ATP level in *L. major* promastigotes was measured using firefly luciferase and its substrate D-luciferin, as described previously [72]. Briefly, 4×10^7^*L. major* cells, grown in low (0.6 mM) glucose medium, were harvested, resuspended in 50 μl of 1X PBS and lysed by sonication. 10 μl whole cell lysate was added to ATP standard reaction solution which was freshly prepared as per manufacturer’s instructions. After incubation for 15 mins at room temperature, luminescence of the sample was measured at 560 nm. ATP concentration for each sample was obtained from the corresponding luminescence value using a calibration curve. The calibration curve was generated by recording luminescence as a function of ATP concentration by measuring the luminescence of increasing concentrations of ATP (75– 600nM). Selection antibiotics were removed from culture medium during the entire course of the experiment.

### Quantification of RNA transcript in *Leishmania*

1 × 10^8^ wild type or mutant *L. major* promastigotes were harvested and total RNA was extracted using TRIzol reagent. Subsequently, DNase I treatment was performed to remove DNA contamination, as per the manufacturer’s instruction. 1 μg of total RNA was used to synthesize cDNA with the help an oligo(dT) primer and Moloney murine leukaemia virus reverse transcriptase (RT). Expression of LmPEPCK (1578 bp), LmCA1 (921 bp), LmCA2 (1887 bp) or LmME (1722 bp) transcript in wild type or mutant *L. major* was checked by semi-quantitative RT-PCR using gene-specific primers P1/P2, P3/P4, P5/P6 or P9/P10 respectively (listed in Table S2). The number of cycles was optimized at 28 after examination of the yield of PCR products at a range of 24–30 cycles. Relative expression of LmPEPCK, LmCA1, LmCA2 or LmME mRNA was normalized using wild type cells as reference sample and rRNA45 gene as an endogenous control. rRNA45 amplification (143 bp) from cDNA was done using primers P13/P14 (listed in Table S2). Relative expression of LmME or LmCA1 gene in wild type cells, grown in high (6.2 mM) or low (0.6 mM) glucose medium, was measured by Real time PCR using primers P15/P16 or P17/P18 (Table S2). Real-time PCR was done on the Step One Real-Time PCR system (Applied Biosystems) using SYBR Green PCR Master Mix. Relative expression level of LmME or LmCA1 mRNA was normalized with wild type cells grown in high glucose medium as a reference sample and rRNA45 gene as the endogenous control using a Comparative C_T_ method as mentioned by the manufacturer.

### Quantification of intracellular parasite load

Infection of J774A.1 murine macrophages with *L. major* was performed as described by us previously [27,28]. Briefly, macrophages were activated with *E. coli* lipopolysaccharide (100 ng/ml) for 6 hrs. Activated macrophages were infected with stationary phase cultures of wild type or overexpressing *L. major* strains, grown in low (0.6 mM) glucose medium, at a parasite to macrophage ratio of 30:1 for 12 hrs. Post-infection, all non-phagocytosed parasites were removed with PBS, and the infected macrophages were incubated for 18 hrs. During this period, wild type *L. major*-infected macrophages were incubated in absence or presence of 25μM ATR7-010 or 0.625μM zineb. Subsequently cells were washed with PBS, fixed with acetone: methanol (1:1) and embedded in anti-fade mounting medium with DAPI. Parasite load (number of amastigotes per 100 macrophages) for each strain/treatment was quantified by counting the total number of DAPI-stained nuclei of macrophages and amastigotes in a field, using an epifluorescence microscope (IX81, Olympus). For each condition, at least 100 macrophages (and corresponding number of amastigotes) were analysed. The percentage of macrophages infected by wild type (untreated or inhibitor-treated) or overexpressing *L. major* strains was determined by counting the total number of DAPI-stained nuclei of uninfected and infected macrophages in a field. For each condition, at least 100 macrophages were analysed.

### Bioinformatic analysis of LmME

LmME (LmjF24.0770) cDNA and protein sequences were obtained from *L. major* gene database [29,30]. Subcellular localization for LmME was predicted by analysing its primary sequence using the online prediction software, TargetP v1.1 to predict presence of any of the N-terminal signal sequence for targeting a protein to ER, mitochondria or chloroplast, TMHMM v2.0 to predict transmembrane helices, BaCelLo to predict presence of a nuclear localization signal, and PTS1 predictor to predict peroxisome targeting signal 1 [73–76].

### Statistical analysis

All statistical analyses were calculated by paired or Student’s t test using GraphPad software. All results were expressed as the mean ± SD from at least 3 independent experiments. P-values indicating statistical significance were grouped into values of ≤0.05 and <0.001; * p≤0.05, ** p<0.01,*** p<0.001.

## Author contributions

DKM, DSP, MA performed the experiments, DKM, DSP analyzed the data and wrote the initial draft of the manuscript, RD conceived and supervised the work, analyzed all data and wrote the final manuscript.

## Acknowledgements

The authors sincerely thank Mr. Ritabrata Ghosh, Mr. Susnata Karmakar, Mr. Kashinath Sahu, and Mr. Sujoy Bose for their technical assistance. The authors are thankful to Drs. Jayasri Das Sarma, Mohit Prasad, and Supratim Datta of IISER Kolkata and Drs. Subrata Adak and Dr. Amogh Sahasrabuddhe of CSIR-IICB and CSIR-CDRI, respectively for their help with various reagents used in this work. The authors also thank Drs. Sankar Maiti and Piyali Mukherjee of IISER Kolkata and Presidency University, respectively, for helpful discussion and constructive suggestions. This research was supported by the Department of Biotechnology (DBT) and Department of Science and Technology (DST) grants BT/PR21170/MED/29/1109/2016 and EMR/2017/004506, respectively. DKM. and DSP were supported by IISER Kolkata fellowships, MA was supported by University Grants Commission fellowship.

## Supplementary materials

**Table S1.** Software-based prediction of subcellular localization of LmME.

| Protein of interest | Software<sup>a</sup> | Analysis report | Inference |
| --- | --- | --- | --- |
| LmME | TargetP v1.1 | Mitochondrial signal peptide is present | Since LmME possesses a mitochondrial signal peptide sequence and has no transmembrane domain, it was predicted to be a mitochondrial matrix protein |
|  | TMHMM v2.0 | Transmembrane domain is absent |  |
|  | BaCelLo | Nuclear localization signal is absent |  |
|  | PTS1 Predictor | Peroxisome targeting signal 1 is absent |  |

<sup>a</sup>Subcellular localization for LmME was predicted by analyzing its primary sequence using online prediction software such as, TargetP v1.1 to predict presence of any of the N-terminal signal sequence for targeting a protein to ER, mitochondria or chloroplast, TMHMM v2.0 to predict transmembrane helices, BaCelLo to predict presence of a nuclear localization signal, and PTS1 predictor to predict peroxisome targeting signal 1.

**Table S2.**
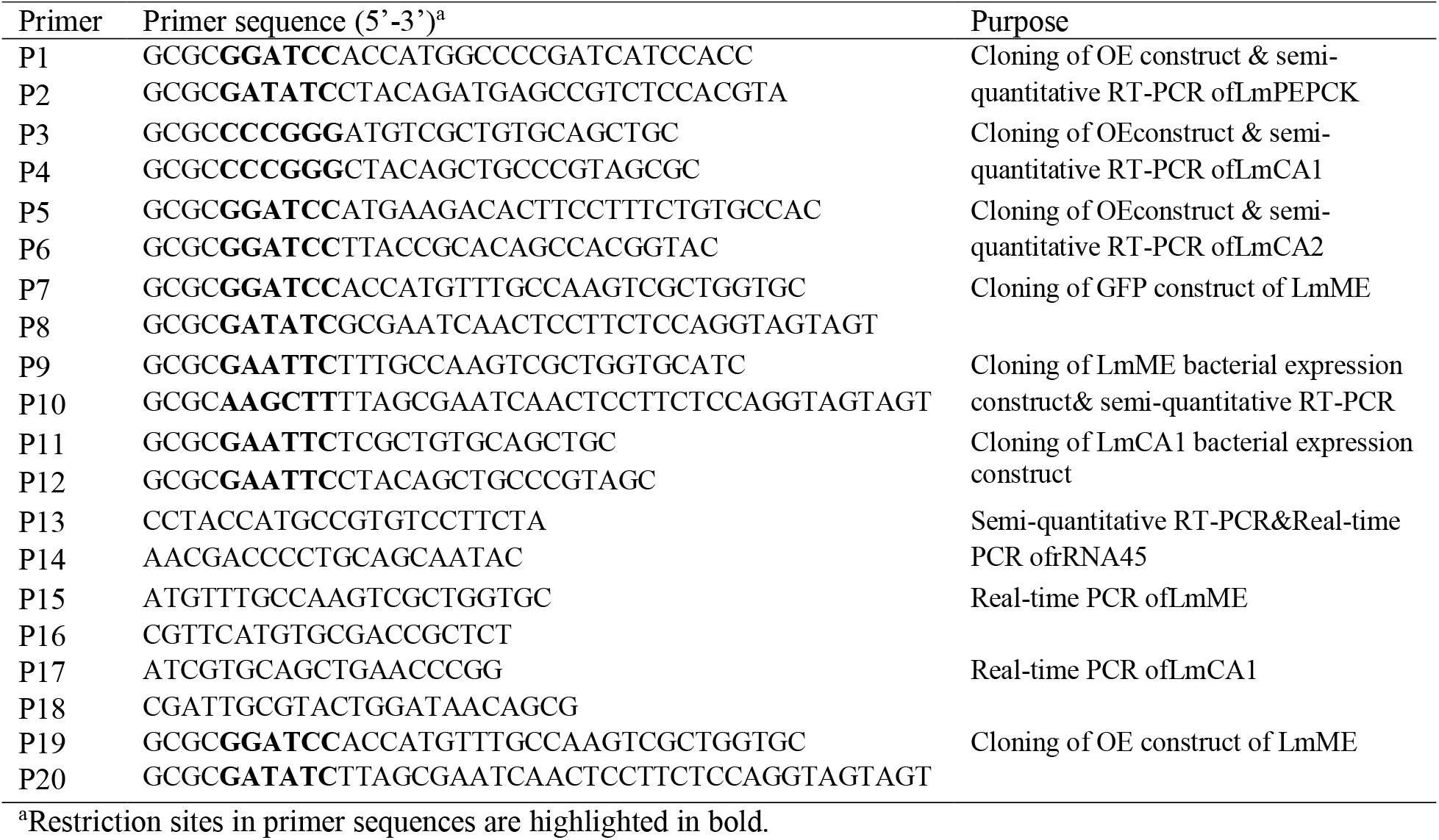
Primers used in this study.

## Legends for supplementary figures

**Figure S1.**
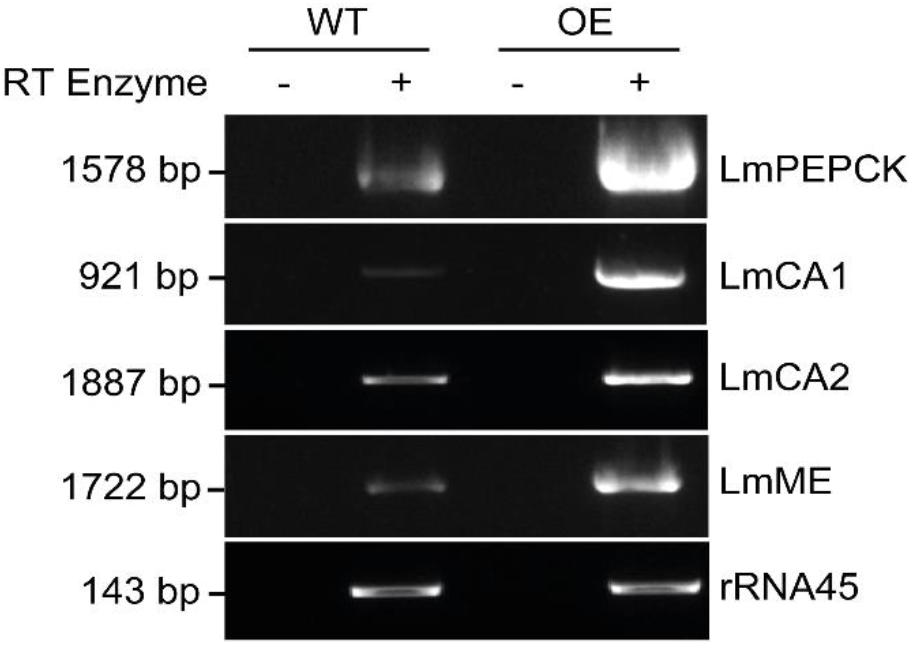
Verification of *L. major* strains overexpressing different genes. Measurement of transcript abundance of LmPEPCK (1578 bp), LmCA1 (921 bp), LmCA2 (1887 bp), or LmME (1722 bp) in wild type (WT) or overexpressing (OE) *L. major* strains by semi-quantitative RT-PCR using primers listed in Table S2 (represented by lanes marked as ‘+’). Respective negative control reactions without RT enzyme are represented by lanes marked as ‘-’. rRNA45 gene (143 bp) was used as the endogenous control.

**Figure S2.**
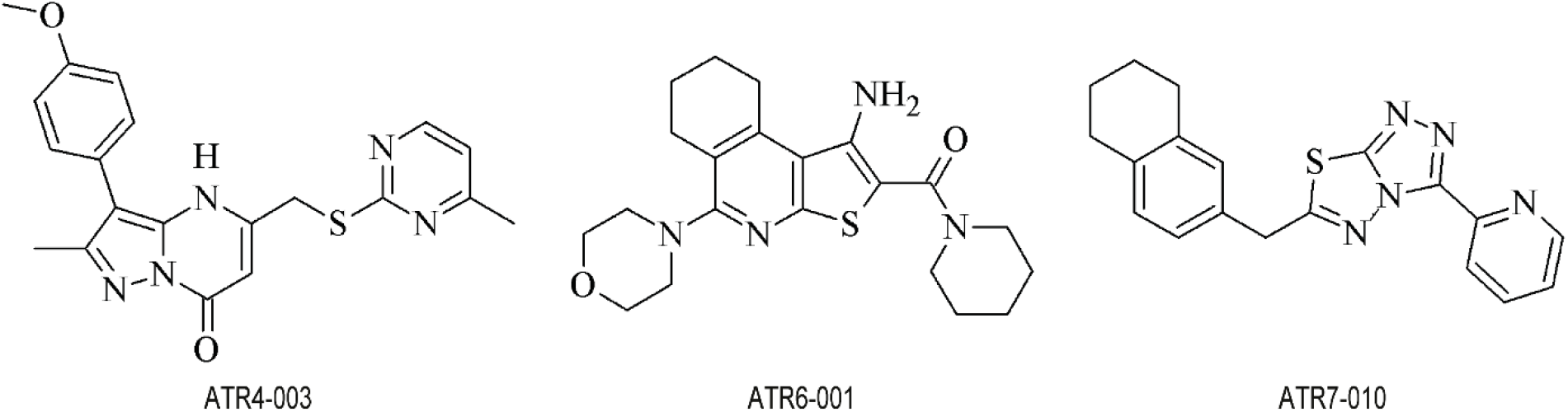
Chemical structures of ME inhibitors used in this study, ATR4-003, ATR6-001, and ATR7-010.

**Figure S3.**
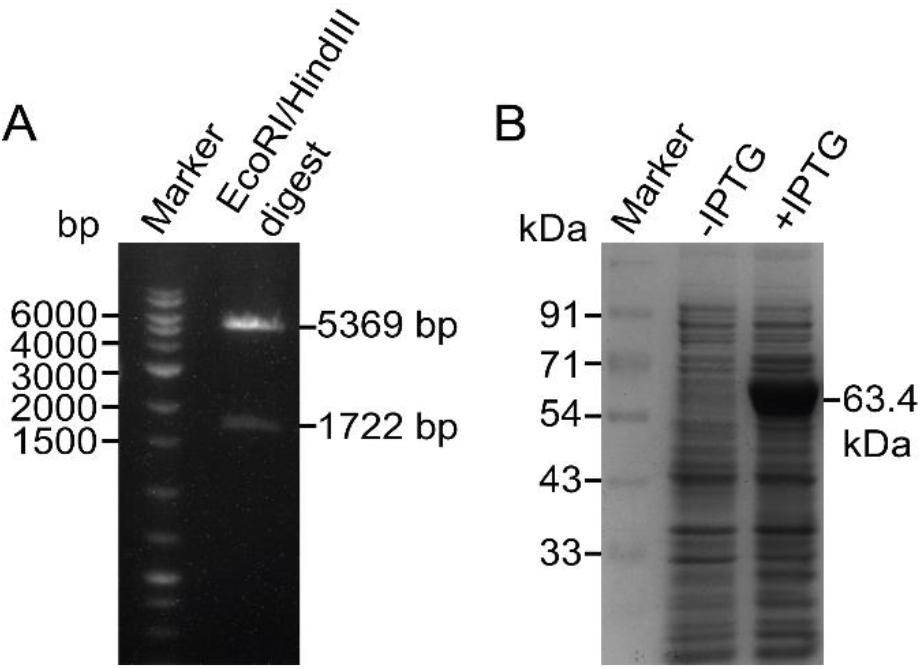
Cloning and expression of 6xHis-tagged LmME in bacterial expression system. (A) Verification of LmME/pET28a+ clone upon restriction digestion with EcoRI and HindIII showing two expected fragments for vector backbone (5369bp) and LmME (1722 bp). (B) Coomassie blue-stained SDS-PAGE showing LmME (63.4 kDa) expression in BL21(DE3) *E. coli* cells grown in presence of 0.5mM IPTG (+IPTG) for 8 hrs at 20°C. LmME was not expressed in bacterial cells grown in absence of IPTG (-IPTG).

**Figure S4.**
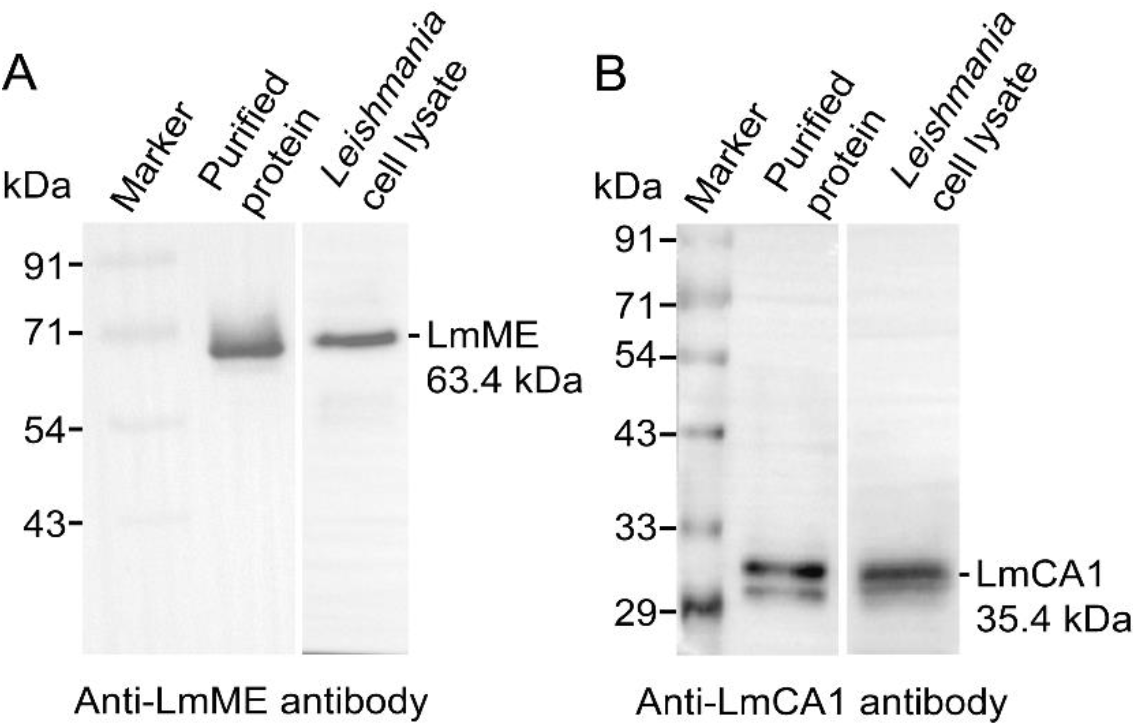
Characterization of LmME or LmCA1 antibodies. (A) 0.5 μg purified LmME protein or 80 μg wild type *L. major* whole cell lysate was subjected to SDS-PAGE and immunostained with anti-LmME antibody (1:4000). LmME band was detected at its predicted molecular weight (63.4 kDa) (B) 9.25 μg purified LmCA1 protein or 120 μg wild type *L. major* whole cell lysate was subjected to SDS-PAGE and immunostained with anti-LmCA1 antibody (1:1000). LmCA1 band was detected at its predicted molecular weight (35.4 kDa).

**Figure S5.**
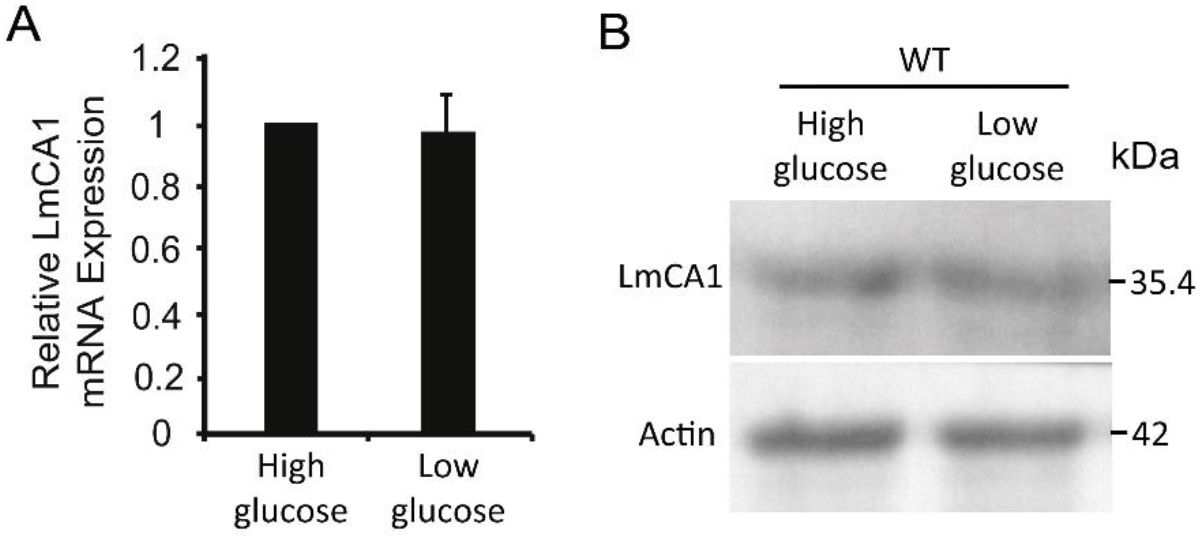
Constitutive expression of LmCA1 in *L. major* cells. (A) Bar graph showing LmCA1 transcript levels in wild type *L. major* promastigotes, grown in high (6.2 mM) or low (0.6 mM) glucose medium for 72 hrs determined by RTqPCR using rRNA45 as endogenous control gene and cells grown in high glucose condition as reference sample. (B) Protein level of LmCA1 (35.4 kDa) in whole cell lysates of wild type (WT) *L. major* promastigotes, grown in high (6.2 mM) or low (0.6 mM) glucose medium for 72 hrs, was checked by western blotting using antibody against LmCA1. Expression of β-actin (42 kDa), detected by anti-β-actin antibody was considered as loading control.

**Figure S6.**
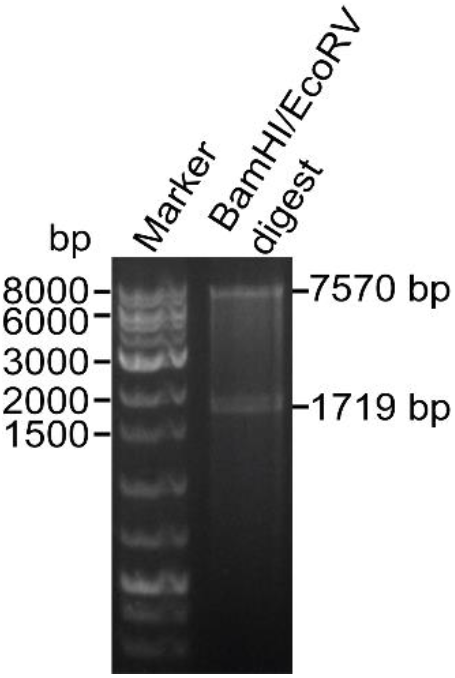
Verification of LmME-GFP clone. Restriction digestion of the LmME-GFP construct with BamHI and EcoRV showing two expected fragments for pXG-GFP vector backbone (7570bp) and LmME without a stop codon (1719 bp).

**Figure S7.**
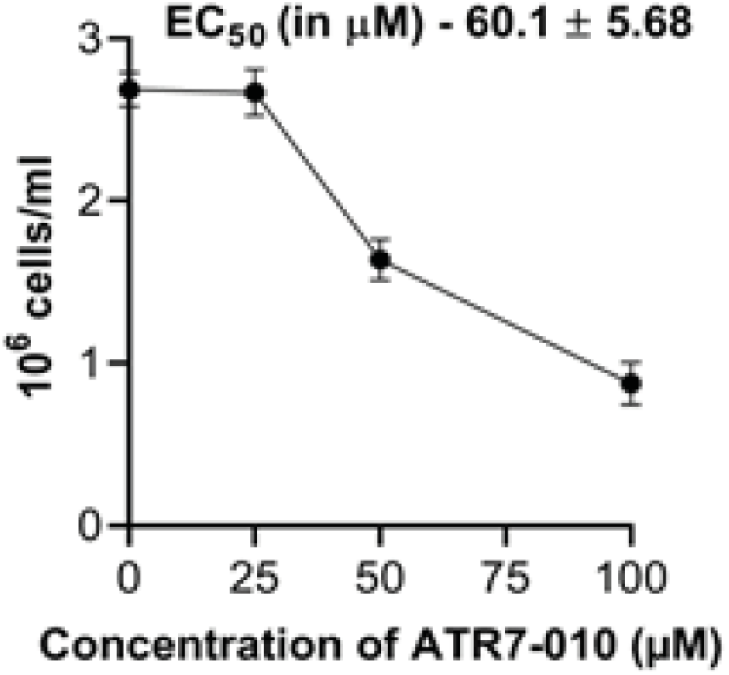
Effect of ATR7-010 treatment on proliferation of J774A.1 macrophage cells. J774A.1 macrophages were grown in absence or presence of indicated concentrations of ATR7-010 and cell growth was measured microscopically after 72 hrs. The EC_50_ value (in μM) of ATR7-010 for J774A.1 cells is given in the index. Error bars represent mean ± SD of values from 3 independent experiments.

